# Substrate recruitment via eIF2γ enhances catalytic efficiency of a holophosphatase that terminates the Integrated Stress Response

**DOI:** 10.1101/2023.12.15.571226

**Authors:** Yahui Yan, Maithili Shetty, Heather P. Harding, Ginto George, Alisa Zyryanova, Katherine Labbé, Amirhossein Mafi, Qi Hao, Carmela Sidrauski, David Ron

**Affiliations:** Cambridge Institute for Medical Research, University of Cambridge, Cambridge CB2 0XY, United Kingdom; Calico Life Sciences LLC, South San Francisco, CA, USA

**Keywords:** Eukaryotic Initiation Factor-2/*metabolism, Phosphorylation, Protein Phosphatase 1/*metabolism, Substrate Specificity, Protein crystallography

## Abstract

Dephosphorylation of pSer51 of the α subunit of translation initiation factor 2 (eIF2α^P^) terminates signalling in the integrated stress response (ISR). A trimeric mammalian holophosphatase comprised of a PP1 catalytic subunit, the conserved C-terminally located ∼70 amino acid core of a substrate-specific regulatory subunit (PPP1R15A/GADD34 or PPP1R15B/CReP) and G-actin (an essential co-factor) efficiently dephosphorylate eIF2α^P^ in vitro. Unlike their viral or invertebrate counterparts, with whom they share the conserved 70 residue core, the mammalian PPP1R15s are large proteins of more than 600 residues. Genetic and cellular observations point to a functional role for regions outside the conserved core of mammalian PPP1R15A in dephosphorylating its natural substrate, the eIF2 trimer. We have combined deep learning technology, all-atom molecular dynamics simulations, X-Ray crystallography and biochemistry to uncover binding of the γ subunit of eIF2 to a short helical peptide repeated four times in the functionally-important N-terminus of human PPP1R15A that extends past its conserved core. Binding entails insertion of Phe and Trp residues that project from one face of an α-helix formed by the conserved repeats of PPP1R15A into a hydrophobic groove exposed on the surface of eIF2γ in the eIF2 trimer. Replacing these conserved Phe and Trp residues with Ala compromises PPP1R15A function in cells and in vitro. These findings suggest mechanisms by which contacts between a distant subunit of eIF2 and elements of PPP1R15A distant to the holophosphatase active site contribute to dephosphorylation of eIF2α^P^ by the core PPP1R15 holophosphatase and to efficient termination of the ISR in mammals.

## Introduction

The integrated stress response (ISR) modifies translation and gene expression to help eukaryotic cells overcome diverse environmental challenges and re-establish cellular homeostasis. Key to the ISR is phosphorylation of eukaryotic translation initiation factor 2 on serine 51 of its alpha subunit (eIF2α), an event triggered by four known kinases (GCN2, PERK, HRI and PKR) responsive to different stresses. Phosphorylated eIF2 (eIF2^P^) inhibits its guanine nucleotide exchange factor, eIF2B. Slow exchange of GDP with GTP in the eIF2γ subunit attenuates translation of most mRNAs and favours translation of specialised mRNAs whose encoded proteins activate a characteristic gene expression program (1–3).

Dephosphorylation of eIF2^P^ terminates the ISR. In mammals, two related regulatory subunits direct the catalytic subunit of protein phosphatase 1 (PP1) to eIF2^P^: PPP1R15B (CReP) is constitutively expressed (4), whilst PPP1R15A (GADD34) is an ISR target gene that is part of a negative feedback loop that quantitatively dominates rates of eIF2^P^ dephosphorylation as cells recover from stress (5–7). PPP1R15s are large proteins, but only ∼70 residues in their C-termini are conserved between the two mammalian proteins and in homologs in other species (8). The conserved core of PPP1R15s forms a stable trimeric complex with PP1 (9) and G-actin (10) and is required for eIF2^P^ dephosphorylation in cells (5, 11, 12). In vitro, the trimer of core PPP1R15, PP1 and G-actin efficiently dephosphorylates its core substrate: the independently-folded N-terminal domain (NTD) of phosphorylated eIF2α (13). Substrate-selective catalytic efficiency of the core holophosphatase is consistent with the structure of a dephosphorylation complex in which G-actin is observed to stabilise an extended segment of PPP1R15A to form a cradle that accommodates eIF2α^P^ NTD and positions pSer51 in the PP1 active site (14).

These phylogenetic and biochemical observation point to the sufficiency of the conserved core of PPP1R15 to serve in eIF2^P^ dephosphorylation and in terminating the ISR. However, in cells, expression of the intact PPP1R15A more potently supresses the ISR than the core fragment (11, 13). Furthermore, there are clues to functionally-important interactions between regions outside the PPP1R15A core and eIF2. On the PPP1R15A side these interactions have been mapped to several short repetitive peptides located N-terminal to the core region, but the interaction has not been mapped on the eIF2 side (15).

eIF2, the physiological substrate of the PPP1R15-containing holophosphatases, is a bi-lobed trimer. The γ subunit, a central component of the larger lobe, interacts with both the β subunit and the C-terminal domain (CTD) of the α subunit which is flexibly attached to the smaller pSer51-containing lobe comprised of the independently folded eIF2α-NTD (16, 17). In yeast, the ISR is terminated by the PP1 (GLC7) catalytic subunit, which is recruited to eIF2 directly by eIF2γ with no intervention by a regulatory subunit (18). The feature enabling direct recruitment of PP1 is missing in mammalian eIF2γ but together these observations hint at the potential importance of additional contacts of the holophosphatase and elements outside the eIF2α substrate lobe of the physiologically-relevant substrate (the eIF2 trimer). Such interactions are missing in experimental systems that contain only the core enzyme and core substrate, but might be mediated by the N-terminal extension of PPP1R15.

Here we have combined deep learning technology, all-atom molecular dynamics simulations and traditional crystallography, biochemistry and cell biology to explore the complex structure of the N-terminal extended PPP1R15A and eIF2 trimer. Our study reveals, in atomic detail, the functionally-important binding of a short helix derived from the PPP1R15A peptide repeats in a hydrophobic pocket on the surface of eIF2γ. These findings extend our understanding of a dephosphorylation event that is key to signalling in mammals and may suggest an unanticipated target for experimental modulation of the ISR.

## Results

### An N-terminally extended PPP1R15A favours dephosphorylation of the eIF2 trimer

To investigate the functional role of elements outside PPP1R15A’s conserved core we purified from bacteria a recombinant fragment of human PPP1R15A^325-636^ that contains the four repeats, previously implicated in eIF2 binding and the core that recruits PP1 and G-actin. This extended PPP1R15A is otherwise wildtype but lacks the hydrophobic N-terminus that associates with membranes and is incompatible with bacterial expression (Fig. 1*A*). In vitro assays demonstrated that dephosphorylation of the eIF2^P^ trimer by the extended PPPR15A^325-636^-PP1-G-actin trimeric holophosphatase was approximately 10 time faster than by the core PPPR15A^533-624^-PP1-G-actin holophosphatase (based on *k*_cat_/*K_m_*, Fig. 1*B* and 1*C*). In contrast, the core holophosphatase dephosphorylated the core substrate (eIF2α-NTD) ∼2 times faster than the extended holophosphatase.

**Fig. 1.**
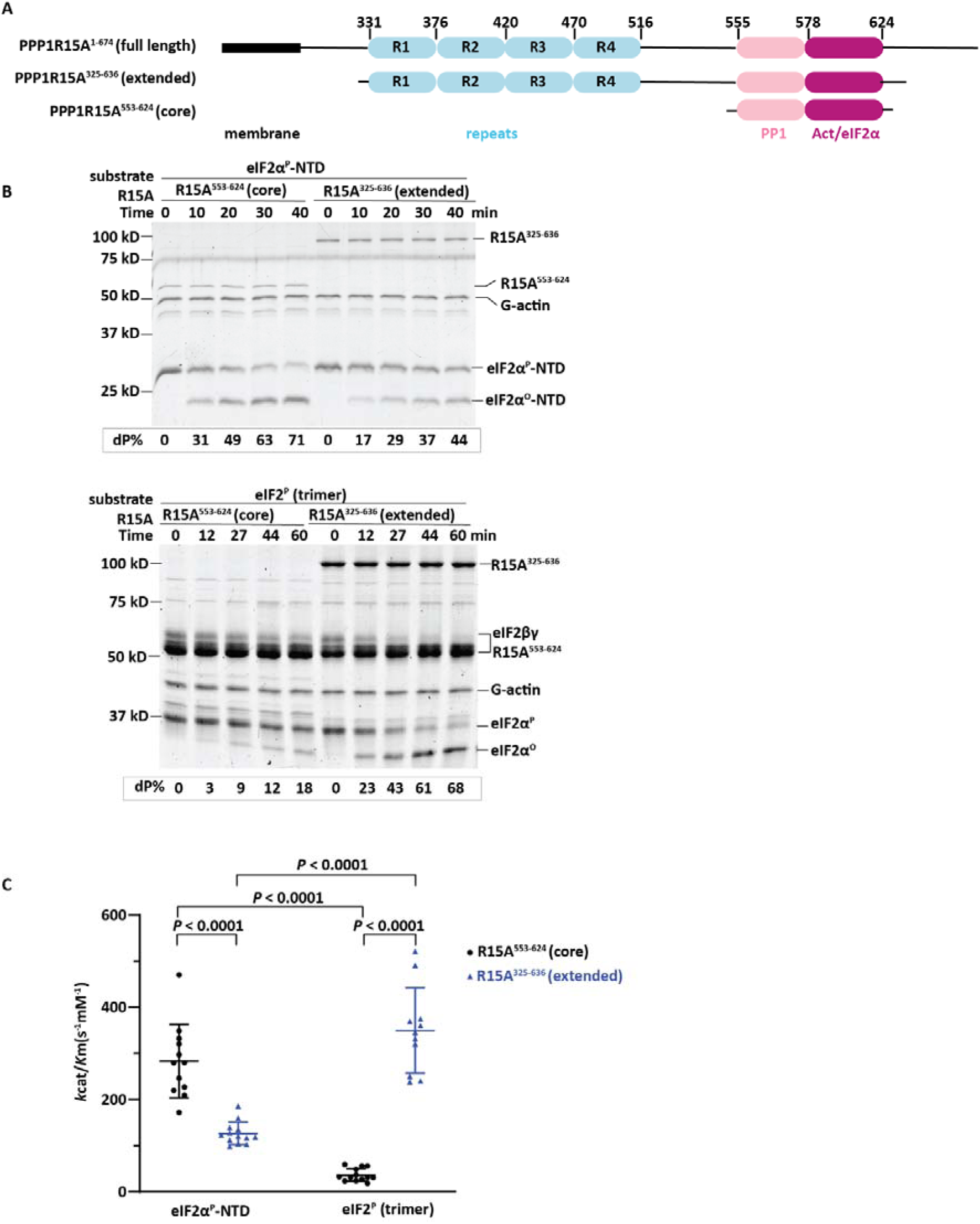
Different substrate preferences of the core and extended PPP1R15A-containing holophosphatase. (A) Schema of human PPP1R15A. Shown is the N-terminal hydrophobic membrane interacting domain, the repeats at the central domain and the C-terminal functional core region that recruits PP1 and G-actin. Elements present in the bacterially-expressed N-terminally extended and core PPP1R15A used experimentally are shown alongside the corresponding residue numbers (based on UniProt O75807) (B) Coomassie stained SDS-PAGE PhosTag gels of dephosphorylation reactions of the core substrate (eIF2α^P^-NTD) or the physiological substrate (eIF2^P^ trimer) by the core holophosphatase (comprised of PPP1R15A^533-624^, G-actin and PP1A) or the extended holophosphatase (comprised of PPP1R15A^325-636^, G-actin and PP1A). Shown is a representative of experiments reproduced three times. (C) Graphic display of the mean ± s.d. of the *k*_cat_/*K*_m_ extracted from all experimental points with the indicated substrates and holophosphatases. *P* values for two tailed parametric t-test are shown.

As previously noted, incorporation of PP1 into a holophosphatase with PPPR15A’s functional core (PPPR15A^533-624^) and G-actin, inhibited non-specific dephosphorylation of glycogen phosphorylase, an irrelevant substrate (Fig. S1) (13). However, incorporation of the catalytic subunit into a PPP1R15A^325-636^ extended holophosphatase was ∼2 times less inhibitory. These observations are consistent with functionally important interactions between the N-terminal extension of PPP1R15A and the eIF2 trimer.

### AlphaFold2-multimer (AFM) and molecular dynamics (MD) simulations predict interactions between PPP1R15A’s repeats and eIF2γ

Our attempts to solve a structure of the extended phosphatase complexed with the eIF2^P^ trimer by cryo-EM were unsuccessful: Two and three-dimensional classification of the particles observed on the grid suggested that many were larger than the core holophosphatase/eIF2α-NTD complex (14). When reconstructed these gave rise to objects of a greater volume than the core enzyme/substrate, but the resolution was too low to plausibly assign different domains of the extended complex to the densities observed (Fig. S2*A*). The NMR spectrum of isolated eIF2α (16) and molecular dynamics simulations of both the eIF2 trimer (Fig. S2*B*) and the extended holophosphatase (Fig. S2*C*) suggested that flexibility between the two lobes of eIF2 and the mostly unstructured nature of N-terminal extension of PPP1R15A may disfavour a fixed orientation of the components of the extended holophosphatase substrate complex, precluding a solution of its structure by CryoEM with the number of particles collected here.

Therefore, we turned to AFM to predict interactions between the eIF2 trimer and PPP1R15A, initially, with the eIF2 trimer and PPP1R15A^325-517^ as inputs. The eIF2 models generated by AFM had high predicted local distance difference test (pLDDT) values for α, C-terminal β, and γ subunits. The predicted aligned errors (PAE) plot of the α subunit (Fig. S3*A*) is explained by the flexible hinge connecting eIF2α-NTD and eIF2α-CTD previously observed in NMR solution structures of eIF2α (PDB 1Q8K) and in the MD simulation (Fig. S2*B*).

Despite a low pLDDT score of PPP1R15A, we noted that three of the top five ranked structures returned by AFM (ipTM+pTM score of 0.59), predicted interactions between a short peptide conserved in PPP1R15A’s repeats and a groove on the surface of eIF2γ. To refine the search process its input was confined to eIF2γ and either one of the PPP1R15A repeats (Fig. 2*A*). High confidence models (ipTM+pTM > 0.7) were returned for repeat 1 (R1, aa. 331-376), 2 (R2, aa.377-420) and 3 (R3, aa. 421-466). In these, the Phe and Trp of a conserved helical ‘<u>F</u>LKA<u>W</u>VY’ motif face into a hydrophobic groove of eIF2γ (Fig. 2*B* and S3*B*). The pLDDT values for residues of the motif were 72-87; and the PAE plot predicted position errors of < 5Å for the motif against eIF2γ. The less well conserved repeat 4 of human PPP1R15A (aa. 471-503) also engaged the same groove of eIF2γ with similar orientation (ipTM+pTM = 0.69) but with a weaker pLDDT score and lower confidence PAE plot (Fig. S3*B*).

**Fig. 2.**
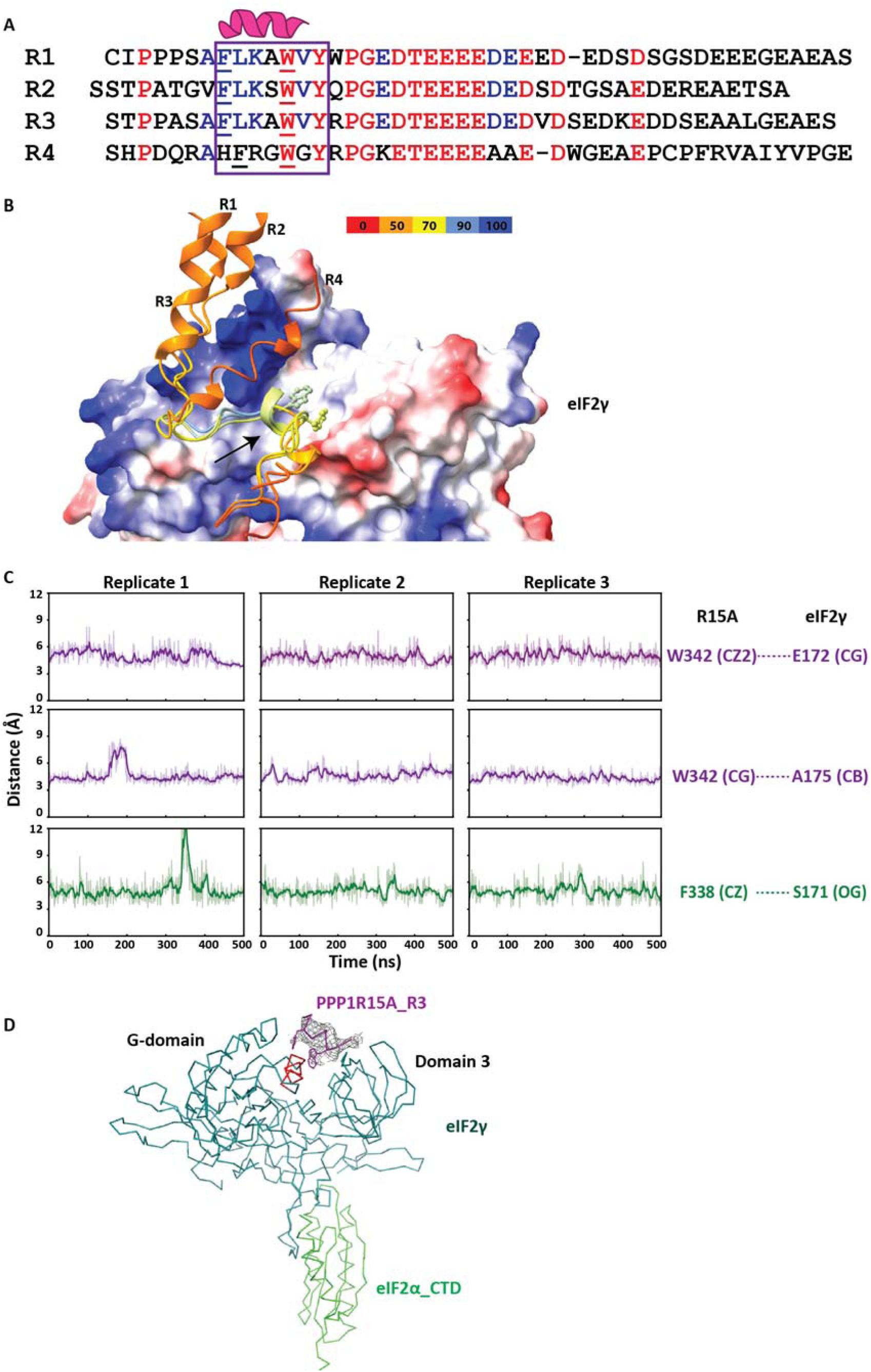
The structural basis of PPP1R15A binding to the γ subunit of eIF2. (A) Alignment of the four repeats region of human PPP1R15A with conserved residues coloured. The helical region that stably interacts with eIF2γ is boxed and the Phe and Trp residues whose sidechains contact eIF2γ are underlined. (B) Overlaid models of individual repeat (R1, R2, R3 or R4) complexed with eIF2γ as predicted by AFM. eIF2γ is shown as electrostatic surface with blue positive charged and red negative charged patches. Repeats peptides are shown as cartoon and coloured by pLDDT (predicted local distance difference test) score. The arrow points at the helix from which the Phe and Trp sidechains face the hydrophobic groove of eIF2γ. (C) Plots of time-dependent variation of distances between the atoms of R15A F^338^ or W^342^ and eIF2γ residues (as indicated) throughout 500 ns of an all-atom MD simulations performed using the AFM predicted complex structure of eIF2γ and repeat 1 (PPP1R15A^331-376^). Shown are three replicates of the simulation. (D) A crystal structure of the eIF2αγ and R3^420-452^ peptide complex as line diagram. The mesh around the R3 peptide is the polder omit map contoured at 3σ, and the red helix in eIF2γ indicates the hydrophobic groove into which the conserved Phe and Trp project their sidechains.

To test the stability of these predicted interactions, all-atom MD simulations of the complex formed by eIF2γ and PPP1R15A repeat 1^331-376^ was performed. Three independent simulations carried over 500 nanoseconds showing stable engagement of the helical ‘<u>F</u>LKA<u>W</u>V’ motif in the eIF2γ groove with the root mean square deviation (RMSD) of 2.4 ± 0.6 Å. The distance between Phe^338^, Leu^339^, Trp^342^, Val^343^ of the helical motif and residues in the eIF2γ hydrophobic groove remained between 3-6 Å throughout the simulation process (Fig. 2*C* and S3*C*). The MD simulations also revealed that although acidic residues conserved in the C-terminal half of the PPP1R15A repeats display considerable flexibility (RMSD=15.6 ± 5.9 Å for three replicates), they still engage in electrostatic interactions with a positively-charged surface of eIF2γ, consistent with rapidly-exchanging salt bridges (Fig. S3*D*). These hydrophobic and charge complementarity interactions result in an overall predicted binding free energy of −11.8 kcal/mol between eIF2γ and PPP1R15A_repeat 1^331-376^.

### A crystal structure of eIF2αγ and PPP1R15A^420-452^

To test experimentally interactions between human eIF2γ and the repeats of PPP1R15A. We sought to crystallise a minimal complex containing eIF2γ and a single repeat. Mammalian expression of eIF2, though suited for the activity assays described above, failed to yield the quantities needed for crystallography and heterogeneous post-translations may hinder the crystallisation. After several attempts, the expression of a multi-cistronic plasmid encoding eIF2αγ and CDC123 in *E.coli* yielded reasonable amounts of monomeric eIF2αγ.

The PPP1R15A^420-452^ (R3) peptide and eIF2αγ dimer, purified separately and combined at 8:1 stoichiometry yielded diffracting crystals and a structure of the complex was determined at 3.35 Å resolution. The interface between the eIF2γ and eIF2α-CTD was well resolved and consistent with previous published eIF2 complexes, whereas no density corresponding to the flexibly-attached eIF2α-NTD was observed. Polder (omit) map unambiguously showed an electron density in the hydrophobic groove between the G-domain and domain 3 of eIF2γ. Residues ^426^SA<u>F</u>LKA<u>W</u>VY^434^ of the PPP1R15A^420-452^ peptide could be built into the density, with the conserved Phe and Trp facing the groove, a configuration in agreement with the AFM predicted model (Fig. 2*D*).

### Phe and Trp on the hydrophobic face of PPP1R15A’s helical repeats contribute to eIF2 dephosphorylation by the extended holophosphatase

A fluorescein-labelled peptide corresponding to repeat 3 (PPP1R15A^420-466^) was titrated with increasing amount of eIF2. This equilibrium binding assay yielded a saturable fluorescence polarisation (FP) signal with a *K*_1/2max_ of 220 nM (Fig. 3*A*). To evaluate the contribution of contacts observed in complexes of eIF2γ and PPP1R15A’s repeat, the Phe and Trp of the helical binding motif of PPP1R15A were mutated to Ala. The fluorescein-labelled double mutant peptide bound the eIF2 trimer weakly and both wildtype and mutant peptides showed similarly low level of unspecific binding with eIF2α-NTD.

**Fig. 3.**
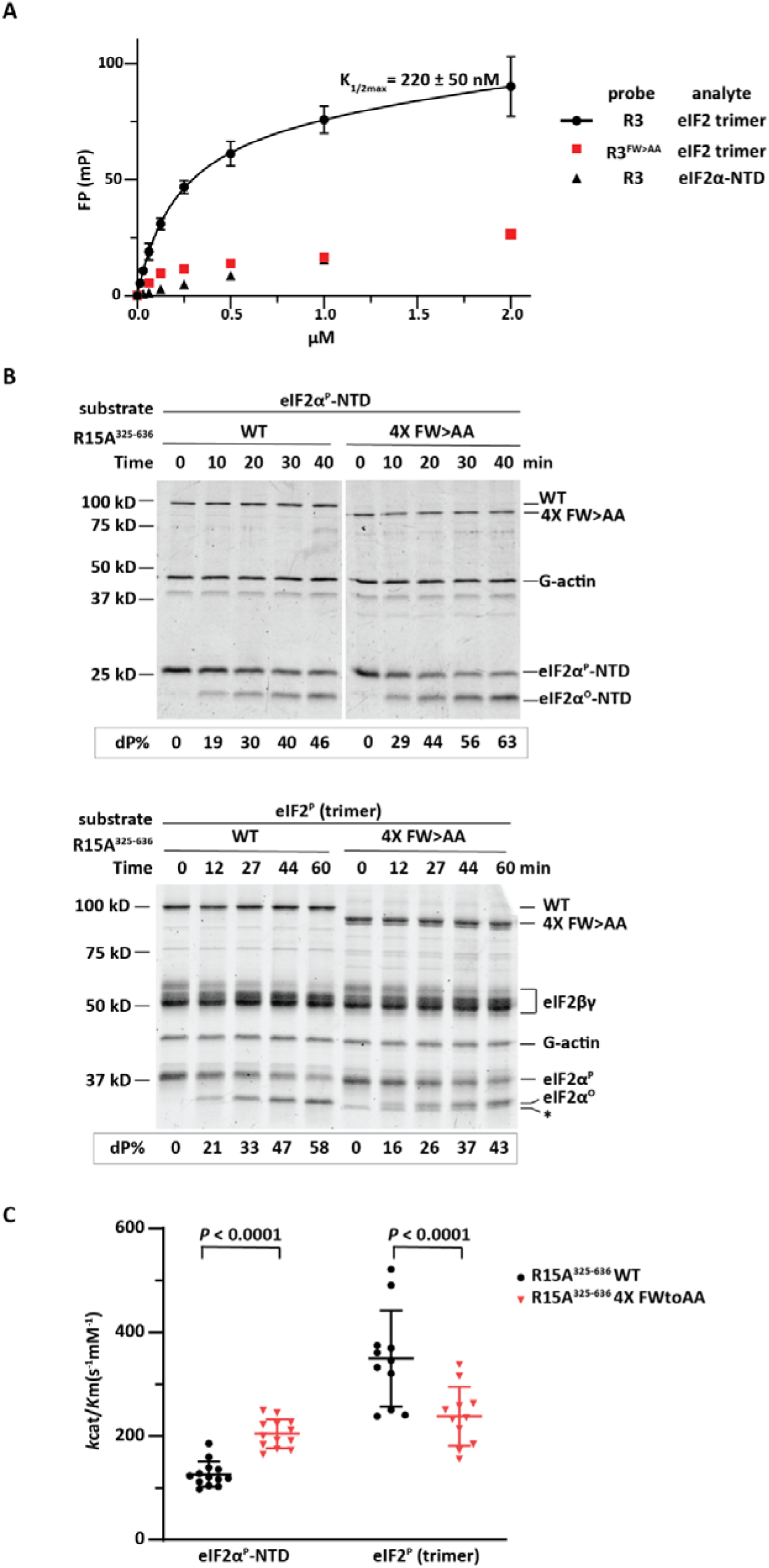
The eIF2γ-contacting Phe and Trp residues of PPP1R15A contribute to eIF2 binding and eIF2^P^ dephosphorylation in vitro. (A) Fluorescence polarization signal as a function of analyte concentrations. Traces from pairings of the wildtype and FW>AA mutant fluorescein labelled R3^420-466^ peptide (as probes) with eIF2 trimer or eIF2α-NTD (an inert reference) are labelled. The binding of the wildtype probe to eIF2 has been fitted to a one site binding model and *K*_1/2 max_ ± 95% confidence intervals is indicated. Shown are the mean ±s.d. from three experiments. (B) Coomassie stained SDS-PAGE PhosTag gels of dephosphorylation reactions of the core substrate (eIF2α^P^-NTD) or the physiological substrate (eIF2^P^ trimer) by a holophosphatase comprised of the extended wildtype or 4X FW>AA mutant (F338A,W342A, F386A, W389A, F428A, W432A, F479A, W482A) PPP1R15A5^325-636^, G-actin and PP1A. Shown is a representative of experiments reproduced three times. (C) Graphic display of the mean ± s.d. of the *k*_cat_/*K*_m_ extracted from all experimental points with the indicated substrates and holophosphatases. *P* values for two tailed parametric t-test are shown.

Next, we replaced these Phe and Trp residues with Ala in all four repeats of the bacterially-expressed extended PPP1R15A^325-636^ (we refer to this mutation as 4X FW>AA) and compared the wildtype and mutant in their ability to dephosphorylate the core substrate, eIF2α^P^-NTD and the physiological substrate, the eIF2^P^ trimer, in the presence of PP1 and G-actin. The 4X FW>AA mutant was selectively impaired in dephosphorylating eIF2^P^ trimer compared with eIF2α^P^-NTD (Fig. 3*B* and *C*). This pattern was also observed in an abbreviated version of the holophosphatase extended with wildtype or mutant version of repeats 3 and 4 (PPP1R15A^420-636^) (Fig. S4). These experiments point to a role for the conserved hydrophobic residues, Phe and Trp, that line one face of the helical repeats of PPP1R15A in binding eIF2γ and suggest that these contacts are functionally important to the dephosphorylation reaction.

### Contacts made by PPP1R15A’s repeats and eIF2 contribute to signal termination in the ISR

PPP1R15A/GADD34 was initially identified as a stress-induced suppressor of the CHOP::GFP ISR reporter gene (5). We revisited this assay to investigate the significance of the interactions uncovered between PPP1R15A’s repeats and eIF2 in the context of PPP1R15A’s capacity to function within this negative feedback loop. Treatment of cells with thapsigargin, an agent that triggers PERK-dependent eIF2 phosphorylation, activated the *CHOP::GFP* reporter (leading to a right-shift of GFP fluorescent signal detected by flow cytometry, Fig 4*A*, top panels). Co-expression of mCherry (by transient transfection) did not affect the CHOP::GFP signal. However, expression of wildtype full length human PPP1R15A (fused with a C-terminal mCherry) repressed the thapsigargin induced CHOP::GFP signal at both low and medium levels of expression (reflected in a shift back to the left of the CHOP::GFP signal in mCherry positive thapsigargin-treated cells). The 4X FW>AA mutant version of the PPP1R15A-mCherry fusion was less effective at suppressing the ISR, and the core PPP1R15A-mCherry fusion was even less effective.

**Fig. 4.**
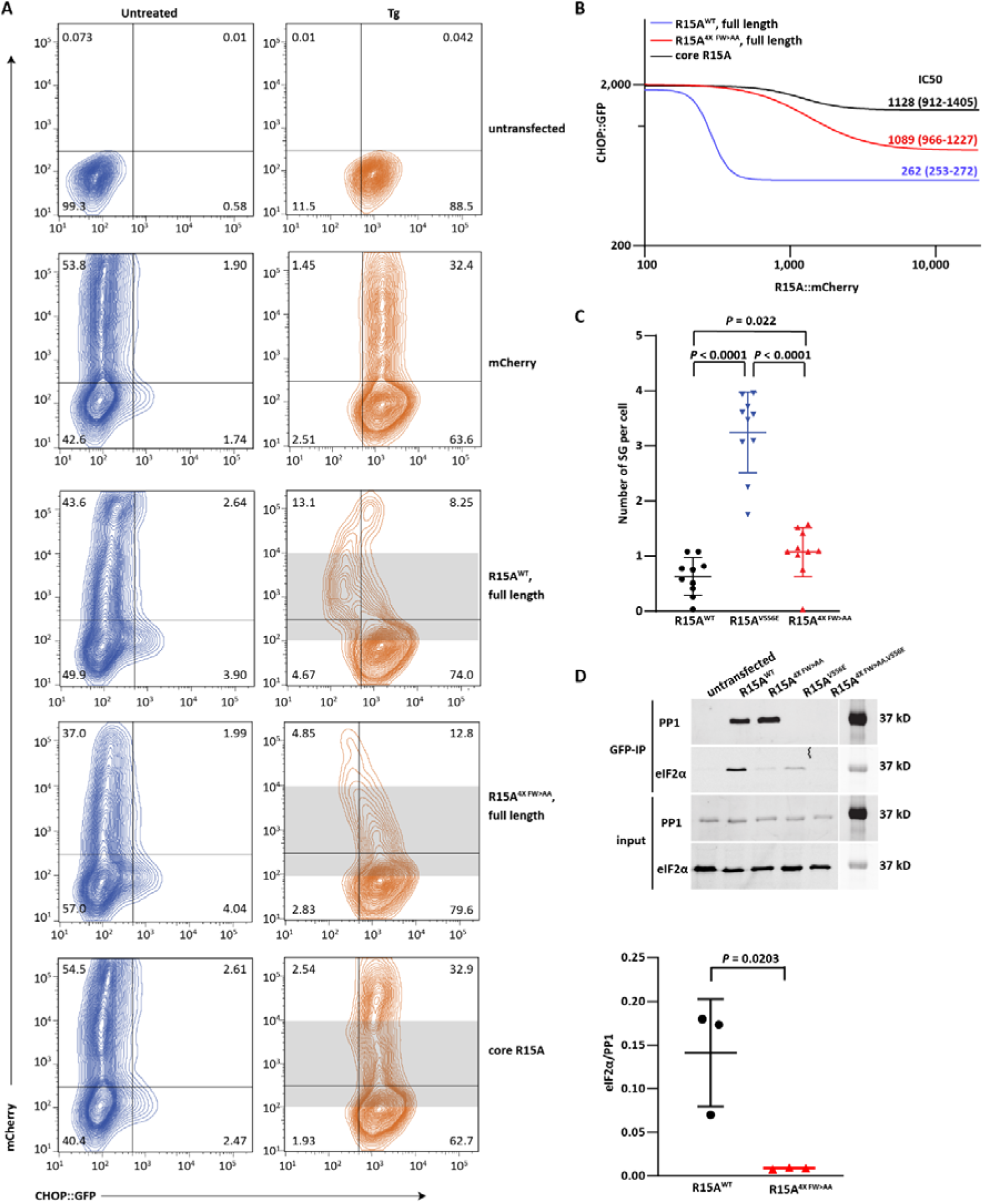
PPP1R15A contacts with eIF2γ contribute to ISR termination in cells. (A) Dual channel flow cytometry plots of untreated and thapsigargin (Tg)-treated CHOP:GFP transgenic CHO-K1 cells transfected with mCherry alone or human PPP1R15A (wild type or mutants as indicated) with mCherry fused to their C-termini. (B) The dual channel fluorescent intensity data from shaded region of plot A was extracted for [inhibitor] (R15A::mCherry) vs response [CHOP::GFP] fitting. IC50 and 95% confidence intervals are indicated for each construct. A combination of fluorescence contamination and protein over-expression artefacts obscure interpretation of signals arising in cells expressing very high levels of PPP1R15A-mCherry, that were therefore excluded from the analysis by setting an upper limit to the shaded zone. (C) Plot of the number of stress granules per arsenite treated cell expressing the indicated PPP1R15A-GFP fusion protein. Shown are all the date points from three experiments performed in triplicate and mean (SD) (*P* values for two tailed parametric t-test are shown). (D) Immunoblots of eIF2α (a marker of the eIF2 trimer) and PP1 (a reference for integrity of the holophosphatase) in the input lysate or recovered in complex with GFP-tagged PPP1R15A (from HEK293T cells transfected with plasmids encoding the indicated wildtype and mutant versions of full-length human PPP1R15A-GFP fusion proteins. Shown is a representative of an experiment reproduced three times. 1% (v/v) of input and 20% (v/v) of immunoprecipitation fractions were analysed. The panel below displays the ratio of the eIF2α/PP1 signal in all three experiments and the mean ± s.d. (*P* values for two tailed parametric t-test are shown).

The mCherry fluorescence signal is a surrogate for the cellular concentration of the different fused PPP1R15A effectors. Therefore the FACS data can be fitted to a nonlinear regression, considering the CHOP::GFP signal as the response and mCherry signal as the inhibitor (in arbitrary fluorescence units). The plateau values of the CHOP::GFP signal supressed by the wildtype PPP1R15A was lower than that of the 4X FW>AA mutant and the IC_50_ (in units of mCherry fluorescence) of the wildtype was 3-4 times lower than the mutant. The core PPP1R15A was even less effective as an ISR inhibitor (Fig 4*B*). Similar observations were also made using wildtype and 4X FW>AA mutant versions of PPP1R15A^325-636^ lacking the N-terminal membrane interacting domain and corresponding closely to the version of the protein used in the in vitro assays, thus aligning the in vitro and in vivo data sets (Fig. S5).

Stress granule formation is another marker of ISR activation (19, 20) that is antagonised by PPP1R15A (14, 21). In PPP1R15A knockout cells (a genetic background sensitised to stress granule formation) introduction of the 4X FW>AA mutant PPP1R15A was less effective than wildtype PPP1R15A in attenuating stress granules abundance (Fig. 4*C*). The eIF2γ-interacting 4X FW>AA mutant PPP1R15A nonetheless retained significant activity, as compared to the V556E mutation that interferes with PP1 recruitment to the holophosphatase. Thus, in both cell-based assays the eIF2γ-interacting mutations have an incomplete but discernible loss-of-function to their ISR-suppressive phenotype.

The 4X FW>AA mutation did not affect the formation of a stable complex between PPP1R15A and the catalytic subunit, as similar amounts of PP1 were recovered in complex with GFP-tagged wildtype and mutant PPP1R15A (immunopurified from transfected cells with a nanobody directed towards GFP). As expected, only background levels of PP1 were recovered in complex with the V556E mutant PPP1R15A. However, considerably less eIF2 was recovered in complex with the 4X FW>AA mutant than the wildtype GFP-tagged PPP1R15A (Fig. 4*D*, in this assay the recovery of eIF2L is deemed a surrogate for the eIF2 trimer). The V556E mutation also greatly decreased eIF2 binding, consistent with the importance of contacts observed between the PP1 active site and eIF2α’s substrate loop in the de-phosphorylation complex (14). The 4X FW>AA mutation further decreased eIF2 binding when introduced in context of the V556E mutant PPP1R15A. Thus, the functional defect in ISR suppression wrought by mutations in PPP1R15A’s Phe and Trp residues that interact with eIF2γ correlated with impaired enzyme-substrate interaction in cells.

## Discussion

Phylogenetics and biochemistry leave little doubt that the ancestral PPP1R15 gene encoded a protein comprised of the conserved 70 residue core that is both necessary and sufficient to form a trimeric phosphatase that selectively dephosphorylates eIF2^P^. Perhaps the best evidence for this is the fact that *Herpes simplex* virus supresses the host anti-viral ISR by directing high level expression of γ34.5, a polypeptide confined to the PPP1R15 core (22, 23). And yet our findings here suggest that the mammalian isoform that accounts for most eIF2^P^ dephosphorylation observed in cells as stress wanes, has acquired a lengthy extension to the core that endows PPP1R15A-containing holophosphatases additional catalytic efficiency when targeting their physiological substrate, the eIF2^P^ trimer.

The biochemical counterpart to this feature of PPP1R15A is an interaction between a shallow hydrophobic groove on the surface of eIF2γ and a repeated segment of PPP1R15A with helical propensity that projects two bulky hydrophobic residues on one side of the helix (Phe and Trp) into the groove. Our crystallographic data is limited to this helix-groove interaction, but MD simulation suggests that PPP1R15A’s affinity for eIF2γ may be buttressed by additional ionic interactions between conserved acidic residues in PPP1R15A and a basic patch adjacent to the hydrophobic groove in eIF2γ. These observations fit with earlier findings pointing to a role for PPP1R15A’s repeats in eIF2 recruitment (15). Mutating the conserved Phe and Trp in PPP1R15A’s repeats compromises ISR suppression in cells and eIF2^P^ dephosphorylation in vitro, suggesting that the contacts established on structural grounds are functionally important. But how do they contribute to catalytic efficiency?

Combined with structures of the core components of the reaction (14), the findings here suggest that the enzyme/substrate complex is comprised of two separate interactions: The eIF2α^P^-NTD (substrate lobe) is engaged by PP1c and the G-actin-stabilised C-terminal extension of PPP1R15’s conserved core, whilst the flexibly-attached eIF2α (C-term)βγ lobe is engaged with one of four PPP1R15A repeats (Fig. 5*A*). Greater affinity of an enzyme for its substrate contributes to catalysis. However, barring an unforeseen rearrangement in eIF2 upon dephosphorylation of eIF2α’s pSer51, the helix-groove interaction discovered here would also feature in the reaction product (dephosphorylated eIF2) and thus limit enzyme turnover. Fine tuning of the kinetics and dominance of contacts made by pSer51 at the PP1 active site, may render the helix-groove interaction a net benefit to catalysis on grounds of enhanced substrate recruitment (Fig. 5*B*), but the gains made via this mechanism might be offset by product inhibition.

**Fig. 5.**
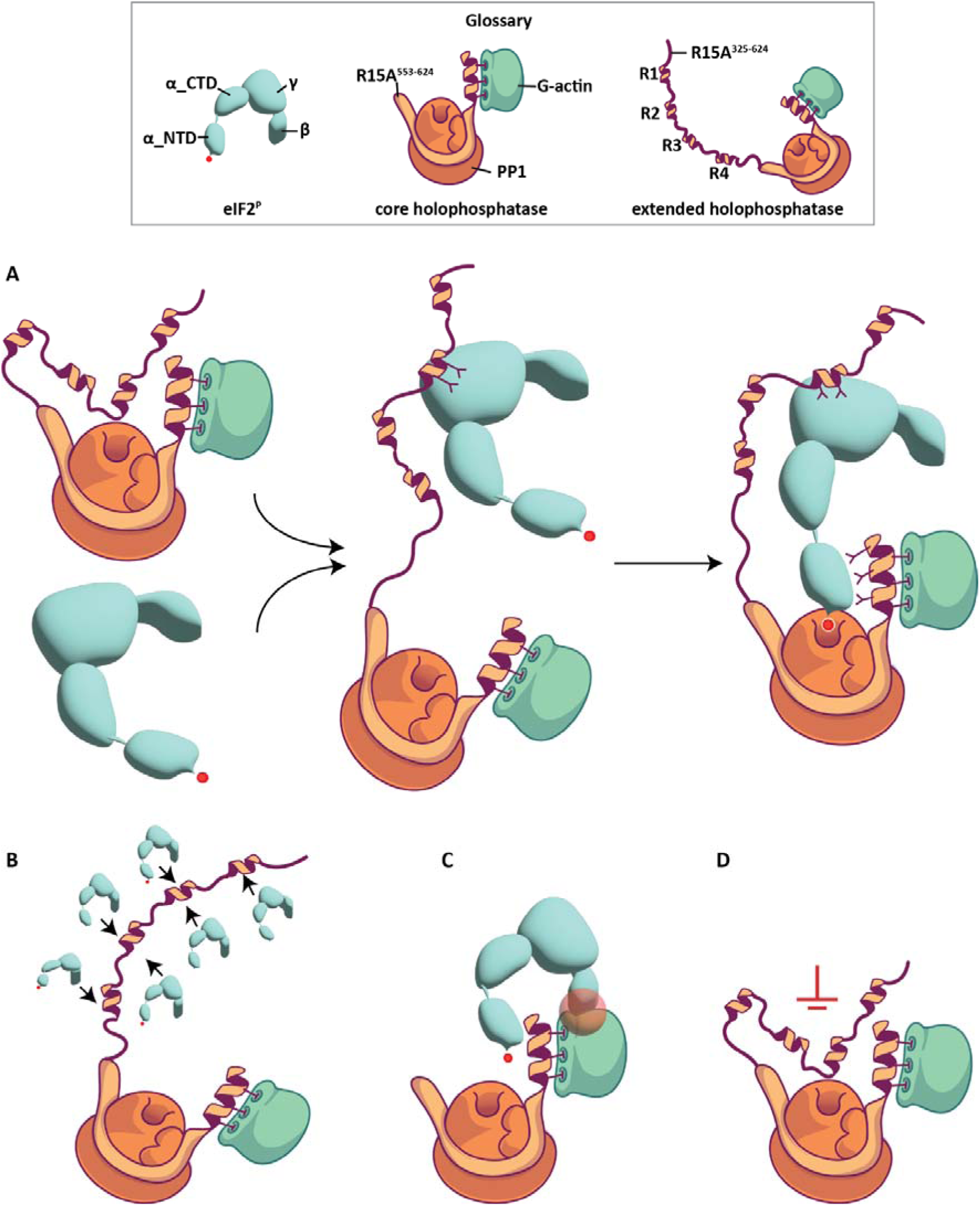
Cartoons depicting hypothetical effects of PPP1R15A-eIF2γ binding on the dephosphorylation reaction. (A) Hypothetical depiction of the formation of an enzyme substrate complex comprised of two sets of interactions: The previously-established contacts made by the eIF2α^P^-NTD and core holophosphatase (14) and the helix-groove interactions between the repeats in N-terminal extension of PPP1R15A and eIF2γ (exemplified here as involving repeat #2). Factors that may affect catalytic efficiency are depicted in the panels that follow. (B) Binding recruits substrates to increase the local concentration of eIF2^P^ in vicinity of the holophosphatase. As the interactions between repeat elements of PPP1R15A and the eIF2γ likely recruit both eIF2^P^ trimer (substrate) and dephosphorylated eIF2 trimer (product). The net benefit of enriched localised eIF2^P^ concentration might be limited by product inhibition. (C) The bulky eIF2 lobe comprised of βγ and α-CTD subunits may interfere with the alignment of the flexibly-attached eIF2α^P^-NTD in the active site (see Fig. S2*B*). Interactions between the extended N-term of PPP1R15A and eIF2γ would stabilise the eIF2α(CTD)βγ lobe limiting an entropic penalty to forming a productive enzyme substrate complex. (D) The N-terminal extended PPP1R15A is largely disordered and may inhibit substrates engagement (see Fig. S2*C*). Helix-groove interactions with eIF2γ would stabilise the N-terminal extension of PPP1R15A limiting another entropic penalty to forming a productive enzyme substrate complex. The box above panel A is a glossary of the reaction’s components (objects adapted from artwork by Claudia Flandoli, http://www.claudiaflandoli.com)

The core holophosphatase is less active in dephosphorylating the eIF2^P^ trimer (the natural substrate) than its isolated eIF2α^P^-NTD lobe (the core substrate), whereas the extended holophosphatase inverts this hierarchy. This suggests that the bulky eIF2α(CTD)βγ lobe interferes with dephosphorylation of the flexibly attached eIF2α^P^-NTD substrate lobe (Fig. 5*C*), interference that may be overcome by contacts between the γ subunit and PPP1R15A’s repeats. According to this model, by engaging the bulky eIF2α(CTD)βγ lobe, the PPP1R15A extension favours docking of the flexibly-attached eIF2α^P^-NTD lobe in the core of the holophosphatase, with the pSer51 loop facing the PP1 active site (Fig. 5*A*).

The extended holophosphatase is less efficient in dephosphorylating the eIF2α^P^-NTD lobe (the core substrate) than the core holophosphatase. This suggests a possibility whereby the eIF2γ-PPP1R15A interaction may also relieve a repressive effect of the PPP1R15A extension on the core holophosphatase or accelerate its assembly into a functional enzyme. The flexibility of PPP1R15A N-terminal extension revealed in the MD simulation could pose an entropic penalty to clearing a path for insertion of the eIF2α^P^-NTD lobe in the enzyme active site (Fig 5*D*). Engagement with eIF2γ might limit this penalty. However, the putative repression relieved by this interaction would have to be substrate dependent, as the PPP1R15A extension does not favour repression of dephosphorylation of phosphorylase A (PYGM^P^), an irrelevant substrate.

PPP1R15 is viewed here through the prism of eIF2^P^ dephosphorylation. This seems justified by the mouse knockout in which sluggish eIF2^P^ dephosphorylation explains much of the phenotype (12). However, others have suggested a role for PPP1R15A’s N-terminal extension in processes unrelated to the ISR (24). Thus, it remains possible that the fitness benefits arising from ISR-unrelated functions of the N-terminal extension of PPP1R15A come at a cost to eIF2^P^ dephosphorylation, a cost that is circumvented by the interactions with eIF2γ.

The hydrophobic groove in eIF2γ is conserved in eukaryotes, whereas the gene duplication that gave rise to PPP1R15A occurred in mammals. Thus, the binding event described here exploited a pre-existing feature of the eukaryotic proteome. This study is restricted to the ISR inducible PPP1R15A subunit; we do not know if functionally-similar features are also found in the constitutive PPP1R15B subunit or in some more distant ancestors. Given the divergence in the non-core regions of PPP1R15A & B (sequence identity < 20%) and the lack of repeats in PPP1R15B it is likely that even if the latter were found to engage eIF2γ it could entail a different process.

Genetics suggests that limiting the rate of eIF2^P^ dephosphorylation may extend the ISR and provide a benefit in certain stress situations (25, 26). This has spawned attempts to access eIF2^P^ dephosphorylation pharmacologically (27, 28). The assays used to identify and characterise putative inhibitors have relied on the dephosphorylation of the core substrate, the eIF2α^P^-NTD lobe, and to date have yielded compounds whose activity as PPP1R15A-selective inhibitors remains contested (29, 30). Here we expand the theoretical target space for inhibitors of eIF2^P^ dephosphorylation and suggest that assays that detect dephosphorylation of the natural substrate, the eIF2^P^ trimer, may offer advantages over those that rely on the core substrate alone in identifying inhibitors that extend the ISR.

## Materials and Methods

### Plasmid construction

PCR-based manipulations, restriction digests, and gene synthesis (Genscript, Piscataway, NJ) were used to mobilize the coding sequence and produces in-frame fusions of the desired proteins with the affinity tags or fluorescent tags, and to create the point mutations indicated in the text. Table S1 lists the plasmids used in this study.

### Protein expression and purification

N-terminal His6-SUMO3 tagged human PPP1R15A (UK2937, UK3133, UK2996, UK3083) with C-terminal fused maltose-binding protein (MBP) were expressed in *E. coli* BL21 T7 Express lysY/Iq cells (New England BioLabs, catalog no. C3013) and purified as previously described, with some modifications (14). Briefly, the supernatant clarified from the cell lysate after overnight induction by 0.5mM isopropylthio-β-D-1-galactopyranoside (IPTG) at 20 °C was applied to pre-equilibrated 5 ml HisTrap column (GE Healthcare). After thorough wash with binding buffer (20 mM Tris–HCl pHL7.4, 0.5 M NaCl, 20 mM imidazole, 0.5 mM TCEP [tris(2-carboxyethyl)phosphine]), the bound protein was eluted with 200 mM imidazole and followed by SENP2 protease (UK2668, at a final concentration of 0.01Lmg/Lml; produced in-house) digestion overnight at 4 °C. The sample was re-applied to the 5 ml HisTrap column to collect the cleaved target protein in the flow through and retain contaminants on the column. Amylose resin (E8021S, NEB) was added to bind MBP-fused target protein. And the bound PPP1R15A protein was eluted with 10 mM maltose after wash. The elution was then applied to HiLoad 16/600 Superdex 200 prep-grade gel filtration column for further purification (buffer: 10 mM Tris–HCl pHL7.4, 0.15 M NaCl and 0.5 mM TCEP). The eluted peak fractions were pooled and snap-frozen in liquid nitrogen and stored at −80 °C.

Human PPP1R15A single repeat peptides (UK2997, UK3098, UK3188) were purified by sequential nickel chelating, SENP2 cleavage and reverse nickel chelating, as described above. After MBP cleavage by TEV protease (UK759, 1:30 w/w ratio, produced in-house) overnight at 4 °C, the sample was purified by anion exchange chromatography (HQ, 1ml, GE Healthcare) eluted with the 0.1-1 M NaCl gradient in 20 mM Tris–HCl pHL7.4 and 0.5 mM TCEP. The eluted peak fractions were applied to HiLoad 16/600 Superdex 75 prep-grade gel filtration column (in 10 mM Tris–HCl pHL7.4, 0.15 M NaCl and 0.5 mM TCEP.) Peak fractions were pooled and snap-frozen in liquid nitrogen and stored at −80 °C.

Human eIF2αγ (UK3136) was purified from *E. coli* lysate by sequential nickel chelating, SENP2 cleavage and reverse nickel chelating, size exclusion chromatography (HiScale S200 increase), TEV cleavage and size exclusion chromatography (HiScale S200 increase), as described above. The eIF2α-NTD (UK2731) was purified as previously described (14).

The eIF2 trimer was recovered from HEK293T cells transiently transfected with plasmids encoding its three subunits, tagged with an N-terminal FLAG followed by a TEV cleavage site. Purification by FLAG-tag affinity chromatography followed by anion exchange chromatography, was performed as previously described (31). Phosphorylation of eIF2 was performed at 30 °C for 2 hours with purified GST-PERK (at 10 nM) in 100 mM NaCl, 25 mM Tris–HCl pHL7.4, 5LmM MgCl_2_, 2.5 mM ATP and 1 mM TCEP. Completion of the reaction was verified by PhosTag SDS–PAGE.

Rabbit PP1A (UK2940, UK2960) was purified by sequential nickel chelating, SENP2 cleavage and reverse nickel chelating, size exclusion chromatography (HiLoad 16/600 Superdex 75 prep-grade), as described above but in presence of 0.5 mM MnCl_2_.

Phosphorylase A (purified from rabbit muscle in the phosphorylated form on serine 15) (PYGM^P^), a non-specific dephosphorylation substrate, was purchased from Sigma (Cat No. P1261)

### Structural analysis

#### Crystallography

7.5 mg/ml eIF2αγ (UK3136) were combined with PPP1R15A_R3^420-452^ (UK3188) at molar ratio of 1:8 in presence of 0.5 mM GMPPNP (ab146659, Abcam) and 5 mM MgCl_2_. A broad screening was done using a Mosquito robot (SPT Labtech) to dispense 100 nl protein with 100 nl of a range of commercial screens to 96-well sitting-drop crystallization trays. And the crystallization took place at 20 °C. The initial small crystals grew at 15% PEG 6000 and 0.1 M Tris-HCl pHL8.5 in ProPlex screen (Molecular Dimensions). After extensive optimisation of pH, PEG concentrations, additive screen and micro-seeding, the best dataset was collected from a single crystal grown in 11-12% PEG 6000, 0.1 M Tris-HCl pHL8.5 supplemented with amino acids microseeded with crushed initial crystals. Diffraction data were collected at beamline i24 at the Diamond Synchrotron Light Source (DLS) at a wavelength of 0.6199 Å and temperature of 100 K using DLS-developed generic data acquisition software (GDA v.9.2). Initial diffraction images were processed by the automatic XIA2 pipeline implementing DIALS (32) for indexing and integration in DLS. Aimless (33) was used for scaling and merging in CCP4i2 (1.1.0) (34). The structure was solved by molecular replacement using Phaser (35) in Phenix (1.20.1-4487) (36) and a copy of AFM predicted human eIF2γ and eIF2α-CTD were found in an asymmetric unit. Although eIF2α-NTD could be accommodated in the crystal packing, it could not be found by Phaser or built manually because of its poor densities. PPP1R15A^426-434^ was manually built to the difference map in Coot (v.0.9.8.7) (37) with reference to the AFM predicted model. Further refinement was performed iteratively using Coot, REFMAC5 (38), phenix-refine (39). MolProbity (40) was consulted for structure validation with 96.1% Ramachandran favored region and no Ramachandran outliers (Table S2). Molecular graphics were generated using UCSF ChimeraX (v.1.6.1) (41) and PyMOL (v.1.3; Schrödinger, LLC).

#### Cryo EM

The complex of PP1^H66K^/R15A^325-636^/G-actin/DNase I/ eIF2α^P^γ purified by HiScale S200 increase size exclusion chromatography was concentrated to 3.5 mg/ml. UltrAuFoil 0.6/1 300-mesh grids (Quantifoil) were glow discharged in residual air at 20 mA for 60 s each side using a Pelco EasiGLOW. 4ul of protein and 0.3ul 3 mM Triton X100 were mixed immediately before deposited on grids and plunged to liquid ethane using a Vitrobot Mark IV (ThermoFisher). Movie stacks were collected on a 300 keV Titan Krios equipped with a K3 camera (Gatan) at the super-resolution counting mode using EPU software. Images were recorded at ×130,000 magnification corresponding to 0.652 Å per pixel using a 20 eV energy filter. Image stacks have 77 frames for an accumulated dose of 53.60Le−/Å^2^ in a total exposure time of 1.04 s. The defocus range was from −2.2 μm to −0.8 μm. A total of 10,576 micrographs were collected from a single grid.

WARP (v.1.0.9) (42) was used for motion correction and CTF estimation, during which the data were binned to given a pixel size of 1.3040 Å. Particles were automatically picked using the BoxNet algorithm of WARP. A total of 418,271 particles stacks generated by WARP were imported to cryoSPARC (v.4.3.0) (43). A few rounds of 2D classification were performed to remove irrelevant particles. A set of 217,482 particles were selected for initial three-dimensional (3D) reconstruction and four initial volumes were generated. Two volumes are too small to fit in the previously determined cryo-EM structure of PP1^D64A^/R15A^553-624^/G-actin/DNase I/ eIF2α^P^-NTD (PDB 7NZM). The third volume has a similar shape to the volume of 7NZM with a correlation of 0.86 and refined to 9.76 Å by non-uniform refinement. The fourth volume is different and larger than 7NZM, but the resolution is too low (9.80 Å by non-uniform refinement) to plausibly assign any models. 2D classification of the particles classified for these two volumes were performed respectively. Considering the flexibility at the hinge connecting eIF2α-NTD and eIF2α-CTD in the full complex, 3D variability analysis was performed for the two large volumes, but did not generate meaningful movements.

#### Structure prediction by AlphaFold-multimer

AlphaFold-multimer(44) was run via the locally installed version of AF2 (versions 2.2.3) (45) on our institutional cluster. All AFM models were generated using default parameters and the prediction was evaluated by the mean of per residue pLDDT (predicted Local Distance Difference Test), PAE (Predicted Aligned Error), PAE-derived pTM (predicted TM) score and the interface pTM score (ipTM). The graphic models were generated using PyMOL (v.1.3; Schrödinger, LLC) and ChimeraX (v.1.6.1).

#### MD simulations

We used AlphaFold-multimer constructs for conducting molecular dynamics simulations. Before initiating the simulations, these deep-learning-derived constructs were refined using the Molecular Operating Environment (MOE) [Inc., C. C. G. Molecular Operating Environment (MOE), 2015.01. 1010 Sherbooke St.West, Suite #910, Montreal, QC, Canada, H3A 2R7 (2015)] QuickPrep module. This preparation included the adjustment of protonation states for all atoms, the capping of both N-and C-termini of the proteins, and the subsequent structure minimization. The minimization process was performed employing the Amber10:EHT force field within the MOE-2022.02 software. During minimization, we imposed MOE default tether restraints on the protein atoms and monitored to maintain the root-mean-square (RMS) gradient of the potential energy below 0.1 kcal mol^−1^Å^−2^.

Subsequently, the refined structures were immersed in a water box and the system was neutralised with an appropriate amount of NaCl to reach a physiological salt concentration of 150 mM. This procedure resulted in systems containing total particles ranging from 237K to 780K atoms. To refine the solvated constructs, each system underwent a 500-step energy minimization using the steepest descents algorithm, as implemented in GROMACS (46). This minimization was succeeded by a canonical ensemble MD simulation, where the system temperature was gradually increased from 0 K to 310 K over 200 ps. This step was followed by a 1 ns MD simulation in an isobaric-isothermal ensemble, maintaining a constant pressure of 1 bar to relax the simulation box for each system.

During these pre-equilibration phases, positional restraints were applied to all heavy atoms with a force constant of 47.8 kcal·mol^-1^Å^-2^, which were gradually reduced to 0 kcal·mol^-1^Å^-2^ in the final equilibration step. The subsequent equilibration runs were conducted in an isobaric-isothermal ensemble as follows:

1. eIF2γ-PPP1R15A^331-376^ with three 500 ns replicates,
2. eIF2 trimer with two 1000 ns replicates,
3. the extended holophosphatase complex, conducted in two replicates (1.3 μs in total)

In our study, we employed the CHARMM36m parameter set (47) to model proteins, phosphotyrosine, ATP, and ions. Water molecules were represented using the CHARMM TIP3P model. The system temperature was maintained at 310 K using the velocity-rescale thermostat (48), which had a damping constant set to 1.0 ps for temperature coupling. Pressure was controlled at 1 bar with the Parrinello-Rahman barostat algorithm (49), applying a 5.0 ps damping constant for pressure coupling. Isotropic pressure coupling was adopted in these calculations.

The Lennard-Jones interactions were truncated at a 12 Å cutoff radius, with a force switch to zero starting at 10 Å. Periodic boundary conditions were implemented in all three dimensions. Long-range electrostatic interactions were calculated using the Particle Mesh Ewald algorithm (50), which utilized a real cutoff radius of 12 Å and a grid spacing of 1.2 Å. The simulation box volume was relaxed using a compressibility of 4.5 × 10-5 bar^-1^. To constrain the water OH bonds, we used the SETTLE algorithm (51) whereas other hydrogen bonds within the system were constrained by the P-LINCS algorithm (52). All MD simulations were performed using GROMACS-2022 (46). For visualization and analysis purposes, Visual Molecular Dynamics (VMD) (53), UCSF Chimera (54), and GROMACS-2022 (46) were employed throughout the manuscript.

#### SDS-PAGE based dephosphorylation assay

The assay followed the procedures described previously (13, 14). The reaction buffer consists of 20 mM Tris pH7.4, 0.1 M NaCl, 0.5 mM MnCl_2_, 1 mM TCEP, 0.02% (v/v) Triton X-100. Rabbit PP1A, PPP1R15A and 300 nM G-actin were incubated for 20 min at room temperature to allow the assembly of the ternary holophosphatase (Fig 1B and 3B: 2.5 nM PP1 and 100 nM R15A for eIF2α^P^-NTD; 1 nM PP1 and 400 nM R15A for eIF2α^P^ trimer. Fig S1: 10 nM PP1 and 400 nM R15A. Fig S3: 1 nM PP1 and 50 nM R15A). The concentration of PP1 and R15A were adjusted to optimally visualise a dynamic range of dephosphorylation over time on gels. The reaction was initiated by adding phosphorylated eIF2α^P^ or PYGM^P^ to an initial concentration of 1 µM.

Equal amount of samples were removed from the reaction at intervals, quenched by SDS sample loading buffer and boiled for 5 min at 70°C, loaded onto 10% or 12.5% SDS–PAGE gel with 50 µM PhosTag reagent and 100 µM MnCl_2_, resolved at 200 V for 60 min or 120 min, stained with Coomassie and scanned by ChemiDoc imaging system (BioRad). The intensity of the phosphorylated and nonphosphorylated bands in each lane was quantified using NIH ImageJ. The percentage of dephosphorylated substrates are indicated for each lane. Reaction rates and *k*_cat_/*K*_m_ were extracted based on (55) as described previously (13, 14) from all the data points of three independent experiments and imported to GraphPad Prism 10 for analysis. *P* values for two tailed parametric t-test were calculated.

#### Binding assay using fluorescence polarization (FP) measurement

Purified human PPP1R15A single repeat peptides (UK2997, UK3098) were labelled with 10-fold molar concentration of fluorescein-5-malamide (16383, Cayman Chemical Company) in the presence of 1 mM TCEP at 4 °C overnight. The free dye was removed by purification using HiLoad 16/600 Superdex 75 prep-grade gel filtration column. Final protein concentration and labelling efficiency was calculated according to the manufacturer’s instructions.

5 ul 100 nM fluorescence probe was mixed with 5 ul of increased concentrations of eIF2 trimer or eIF2α-NTD. The reaction buffer consists of 20 mM Tris pH7.4, 0.1 M NaCl, 1 mM TCEP, 0.01% (v/v) Triton X-100. The FP signal was measured using CLARIOstar (BMG Labtech) with the excitation wavelength at 482 nM and emission wavelength at 530 nM. The gain was adjusted using

the well with the probe alone. The FP signal was plotted against the eIF2 concentration and fitted for one site binding in GraphPad Prism 10. Three independent experiments were performed.

### ISR activity in cells

#### Monitoring ISR in CHOP::GFP report cell line by flow cytometry

The ISR reporter gene CHOP is C-terminally fused with the green fluorescent protein coding sequence (CHOP::GFP) and incorporated stably as a transgene in CHO cells (5). CHO cells were maintained in Ham’s F12 medium (Sigma) supplement with 10% fetalclone II serum (HyClone) and 1X antibiotic mix (penicillin-streptomycin) (Sigma) and 2 mM L-glutamine (Sigma). Cells were cultured at 37°C in 5% CO_2_.

For flow cytometry, cells were seeded in 6-well plates at a density of 1.5 x 10^5^. 24 hours later cells were transfected with 2 ug DNA (mCherry tagged PPP1R15A DNA and empty carrier DNA at a ratio of 1:19) using the lipofectamine LTX system (Life Technologies) at a ratio of 3 μl lipofectamine LTX to 1 μl Plus reagent for every nanogram of DNA in 200 μl of Opti-MEM (Thermo Fisher Scientific). At 12 hrs post transfection cells were treated with 0.5 μM thapsigargin (Calbiochem) to activate the CHOP::GFP reporter. At 24 hours post-transfection cells were prepared for flow cytometry analysis. Cells were washed twice in chilled PBS-1 mM EDTA and released by incubating in PBS-4 mM EDTA for 5 mins at room temperature. Samples were transferred into flow cytometry tubes placed on ice and analysed within 2 hours.

Flow cytometry analysis was performed using LSRFortessa cell analyser (BD Biosciences). Acquisition was conducted for GFP (excitation laser 488nm, filter 530/30; voltage) and mCherry (excitation laser 561, filter 610/20; voltage). Gating strategy described in supplementary data. Flow cytometry data was acquired using FACSDIVA (v.8.0.1 BD Bioscience) and analysed using FlowJo v.8.0. CHOP::GFP and mCherry values correlated to mCherry values in the range of 10^2^-10^4^ were subsequently transferred as csv (scale values) to GraphPad Prism V10 to perform relevant statistical analysis. [Inhibitor] vs response (variable slope-4 parameters; least squares regression with no special handling of outliers) curves were plotted to extract IC_50_ values and 95% confidence intervals. Three independent experiments were reproduced.

#### Stress granules formation

Stress granules formation experiments were modified from the previous protocols (20). U2OS cells were cultured in Dulbecco’s modified Eagle’s medium (DMEM, Gibco) supplemented with 10% fetal bovine serum (FBS, Sigma) and 1X antibiotic-antimycotic solution (Gibco). Cells were housed in an incubator at 37°C, 5% CO_2_, 20% O_2_ and passaged every 2-3 days with trypsin.

Cells were plated at 0.3 x 10^6^ per well in 6-well plates and transfected the next day with 100 ng GADD34-GFP or control vector using 4 µl Lipofectamine 2000 (Thermo Fisher Scientific) following the manufacturer’s instructions and allowed to recover overnight. Transfected cells were trypsinized and 0.02 x 10^6^ cells were plated per well in CellCarrier 96 well plates (Perkin Elmer). The next day, cells were treated with 100 µM sodium arsenite (Merck, cat # 106277) for 1 hour before fixing in 4% paraformaldehyde for 15 minutes, washing 3x in PBS and blocking in block buffer (PBS, 5% FBS, 0.5% Triton X-100) for 1 hour. Anti-G3BP1 (BD) diluted (1:2000) in labelling buffer (PBS, 0.1% Tween, 5% FBS) was applied and incubated at 4°C overnight. Following 3 PBS washes, Alexa Fluor 495 labelled secondary antibody (Thermo Fisher Scientific) was applied (1:10000) in labelling buffer for 1 hour, followed by 3 PBS washes, staining with DAPI (1:5000) in PBS for 15 minutes and 3 PBS washes. All labeling steps were performed at room temperature and samples were protected from light throughout.

Cells were imaged in fresh PBS. Fluorescent imaging was performed on the Opera Phenix High Content Screening system (Perkin Elmer) fitted with a 40X water objective. Nucleus, cell, and spot segmentation were performed using Harmony software (Perkin Elmer). For quantification of stress granule content, 9 random fields were selected and a total of >100 cells were scored across 3 wells per condition for each of 3 independent experiments. Data plotting and one-way ANOVAs were performed on GraphPad Prism software.

#### PPP1R15A pull-down assay

HEK293T cells cultured in DMEM with 10% Fetal Calf Serum were transfected using linear polyethylenimine (MW ∼25,000, Cat#23966 Polysciences Inc., Warrington, PA), 40 µg/ 10 cm dish and plasmid DNA, 10 µg/ 10 cm dish. Thirty-six hours later, complexes containing GFP-tagged PPP1R15A were recovered on a resin coated with anti-GFP nanobodies (GFP-Trap, Proteintech, Planegg, Germany), following lysis in the manufacturer’s recommended buffer supplemented with 100 µM TCEP. Proteins were eluted from the resin in 1% SDS; 50 mM DTT, resolved by 12% SDS PAGE and blotted onto a PVDF membrane, that was subsequently probed with a rabbit polyclonal serum to human eIF2α-NTD at 1:3,000 (13) and detected with an IR800 secondary Goat anti-rabbit antiserum (Li-Cor Cat. # 926–32211). The blot was subsequently probed with a mouse monoclonal to protein phosphatase 1 catalytic subunit (56) and detected with an IR680 secondary Goat anti-mouse antiserum (Li-Cor Cat. # 926–68070), on a Li-Cor Odyssey scanner (Li-Cor, Lincoln, NE).

#### Data, Materials, and Software Availability

The atomic coordinates and structure factors of the eIF2αγ/R15A^420-452^ complex structure have been deposited to Protein Data Bank with accession code of PDB 8QZZ. Plasmids, key reagent resources and software are included in the manuscript and supplement information (Table S1 and S3).

## Acknowledgements

We thank the Huntington lab for access to the Octet machine, Reiner Schulte and Gabriela Grondys-Kotarba from the CIMR flow cytometry core facility team for help with cell sorting, Alexander Faille (CIMR) for advice on CryoEM, Graham Pavitt (University of Manchester) and Takuhiro Ito (RIKEN) for advice on eIF2 expression in bacteria, Lucy Dalton and Stefan Marciniak (CIMR) for sharing mammalian expression vectors of GFP-tagged human GADD34, Stefan Marciniak for comments on the manuscript and Claudia Flandoli (http://www.claudiaflandoli.com) for artwork. Diamond Light Source i24 (mx28677-33) for X-ray crystal structure data collection. S. Hardwick, D. Chirgadze and L. Cooper (Cryo-EM facility, Department of Biochemistry, University of Cambridge) for Cryo-EM data collection. This work was supported by Wellcome Trust Principal Research Fellowship to DR (Wellcome 224407/Z/21Z).

## Author contributions

YY: Designed research, performed research, analyzed data, wrote the paper

MS: Performed research, analyzed data

HPH: Designed research, analyzed data

GG: Contributed new reagents or analytic tools

AZ: Contributed new reagents or analytic tools

KL: Performed research, analyzed data

AM: Designed research, performed research, analyzed data

QH: Designed research, analyzed data

CS: Designed research, analyzed data

DR: Designed research, performed research, analyzed data, wrote the paper

**The authors declare no competing interests.**

---

**Fig. S1.**
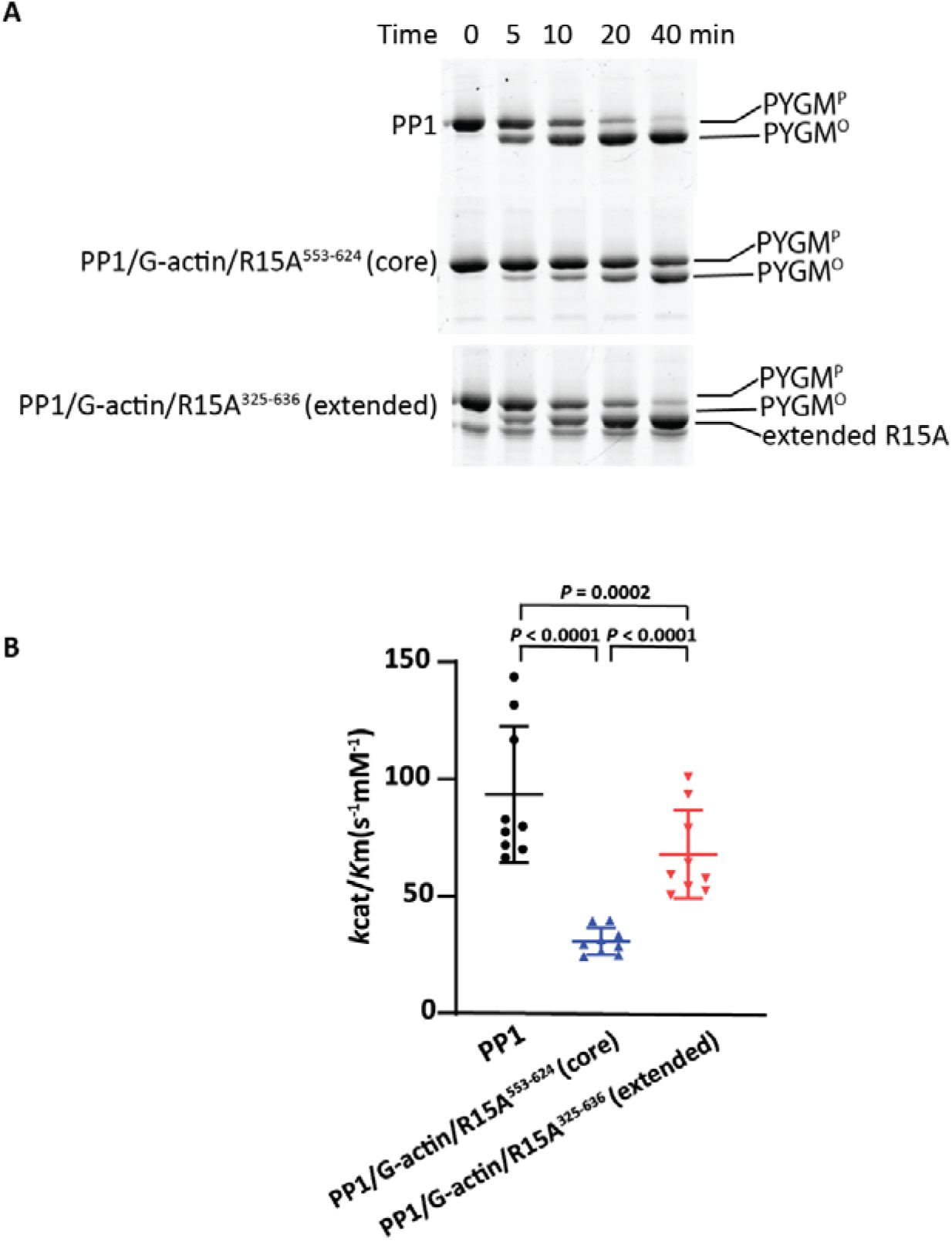
Dephosphorylation of a non-specific reference substrate [glycogen phosphorylase A (PYGM)] by apo PP1, PPP1R15A core and extended holophosphatases. (A) Coomassie stained SDS-PAGE PhosTag gels of PYGM^P^ following dephosphorylation in vitro by apo PP1, core or extended holophosphatases. 10 nM PP1, 300 nM G-actin and 400 nM core R15A or extended R15A were incubated for 20 min at room temperature, then mixed with 1uM PYGM^P^ for different time intervals at 30°C. Shown is a representative of experiments reproduced three times. (B) The panel displays the mean ± s.d. of the *k*_cat_/*K*_m_ extracted from three independent experiments with the indicated holophosphatases. *P* values for two tailed parametric t-test are shown.

**Fig. S2.**
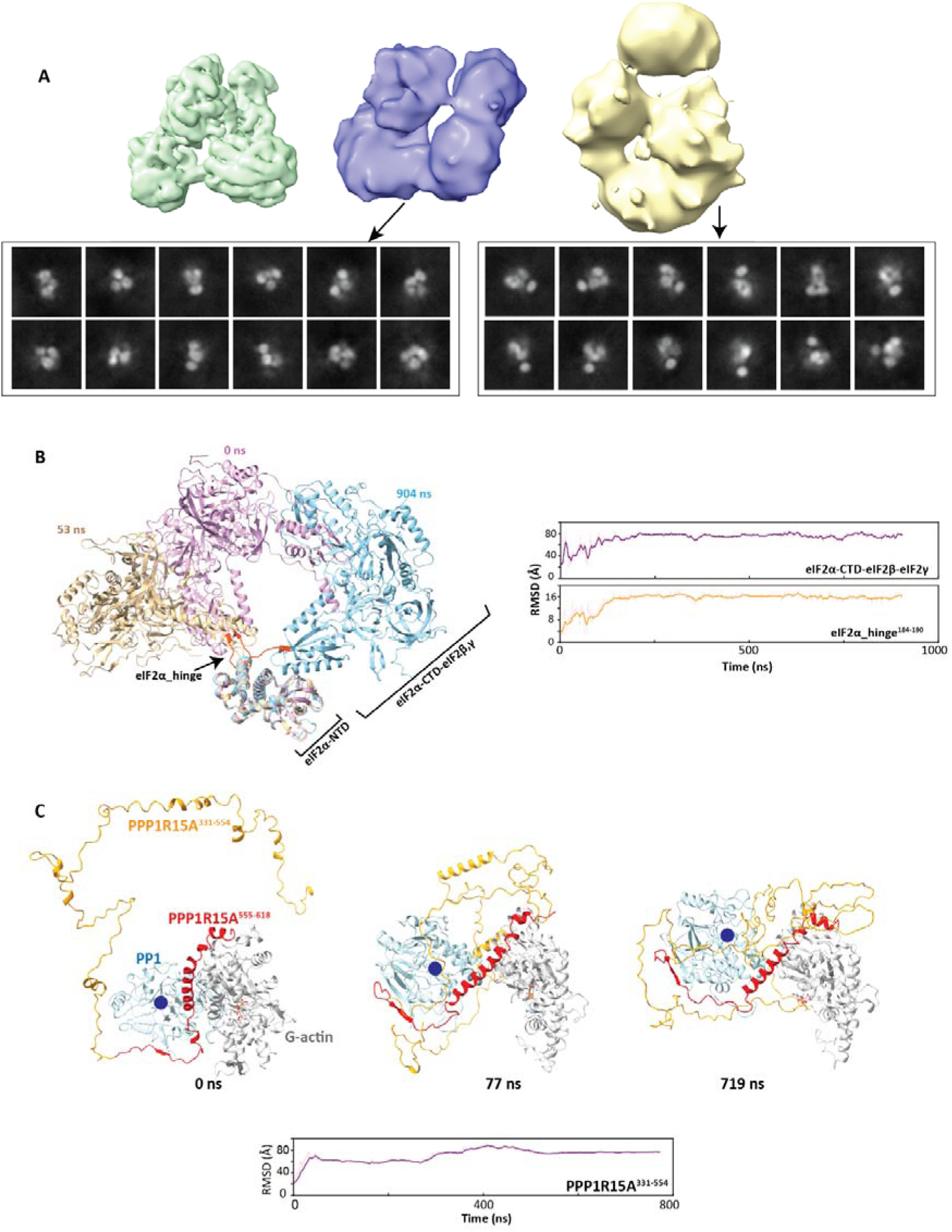
Flexibility of eIF2 trimer and the N-terminally extended PPP1R15A-containing holophosphatase hint at the difficulty in solving the structure of a dephosphorylation complex of eIF2^P^. (A) Two and three dimensional reconstructed images of complexes of eIF2^P^ and an extended holophosphatase comprised of human PPP1R15A^325-636^, PP1A and G-actin. The green volume on the left represents the previously determined cryo-EM structure of PP1^D64A^/G-actin/DNase I/R15A^553-624^/eIF2α^P^-NTD (PDB 7NZM). Two low resolution cryo-EM volumes were obtained from the complex of PP1^H66K^/G-actin/DNase I/R15A^325-636^/eIF2α^P^γ (blue and yellow surface contoured at a similar level) by ab-initio reconstruction in CryoSPARC: the blue is similar to the 7NZM (map correlation is 0.86) and the yellow volume is different and bigger than the former. Below are selected 2D classes generated from the particles classified for these two volumes. Despite the presence on the grid of particles of a size consistent with the eIF2^P^ dephosphorylation complex, we were unable to solve the structure at a meaningful resolution. (B) MD simulations of the eIF2 trimer Shown is a simulation starting from the highest ranked AFM model of eIF2. The left panel display overlaid three distinct conformations of the eIF2 trimer throughout the simulation trajectory, underscoring the flexibility of the eIF2α-CTD, eIF2β, and eIF2γ, compared to the relatively stable eIF2α-NTD. The right panels depict the RMSD plots for eIF2α-hinge (residue 184-190) and the composite eIF2α-CTD-eIF2β-eIF2γ. RMSD calculations were performed after aligning the entire trajectory on the eIF2α-NTD, because of its considerable stability (RMSD 2.3 ± 0.3 Å). (C) MD simulations of the extended holophosphatase MD simulations of the extended holophosphatase highlight the notable flexibility of the N-terminal extension of PPP1R15A (PPP1R15A^331-354^). Three specific conformations of the extended holophosphatase are displayed over the course of the simulation trajectory. The components are coloured as indicated and the blue circle represents the PP1 active site. RMSD of PPP1R15A was calculated with respect to PP1 and G-actin proteins. RMSD values reported and computed were specifically for the CL atoms. Solid lines in plot panels represent the exponential moving average throughout the MD simulations for each respective replicate.

**Fig. S3.**
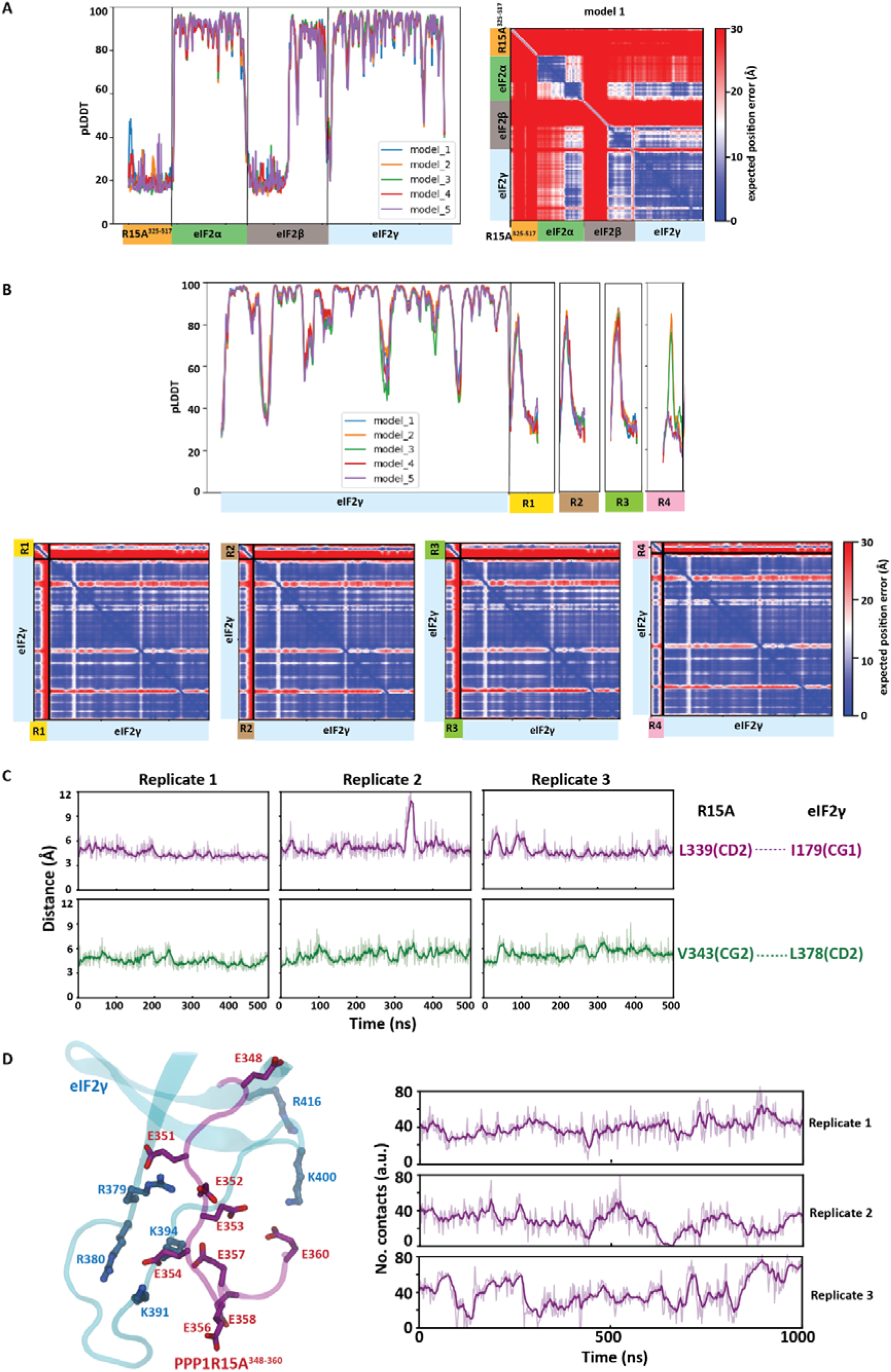
AFM prediction and MD simulations of eIF2γ and PPP1R15A complexes. (A) The pLDDT (predicted local distance difference test) and PAE (Predicted Aligned Error) plots for AFM predictions of the human eIF2 trimer in complex with the repeat-containing region of PPP1R15A^325-517^. (B) The pLDDT and PAE plots for AFM prediction of complexes of human eIF2γ and individual repeats of PPP1R15A: R1^331-376^, R2^377-420^, R3^421-466^ or R4^471-503^ respectively. (C) Plots of time-dependent variation of distances between the atoms of human PPP1R15A L^338^ or V^343^ and the indicated residues of human eIFγ throughout 500 ns of an all-atom MD simulations performed using the AFM predicted complex structure of eIF2γ and repeat 1 (PPP1R15A^331-376^). Shown are three replicates of the simulation. (D) The dynamic interactions between the acidic segment of PPP1R15A’s repeat 1 region (PPP1R15A^348-360^) and the positively charged surface of eIF2γ. The left panel displays the interface interaction captured at time=500 ns from the calculations of replicate 1. The right panel presents a stability analysis based on the count of contacts between PPP1R15A^348-360^ and the positively charged surface of eIF2γ throughout the MD simulations. The cutoff radius for computing the contact was set at 6 Å.

**Fig. S4.**
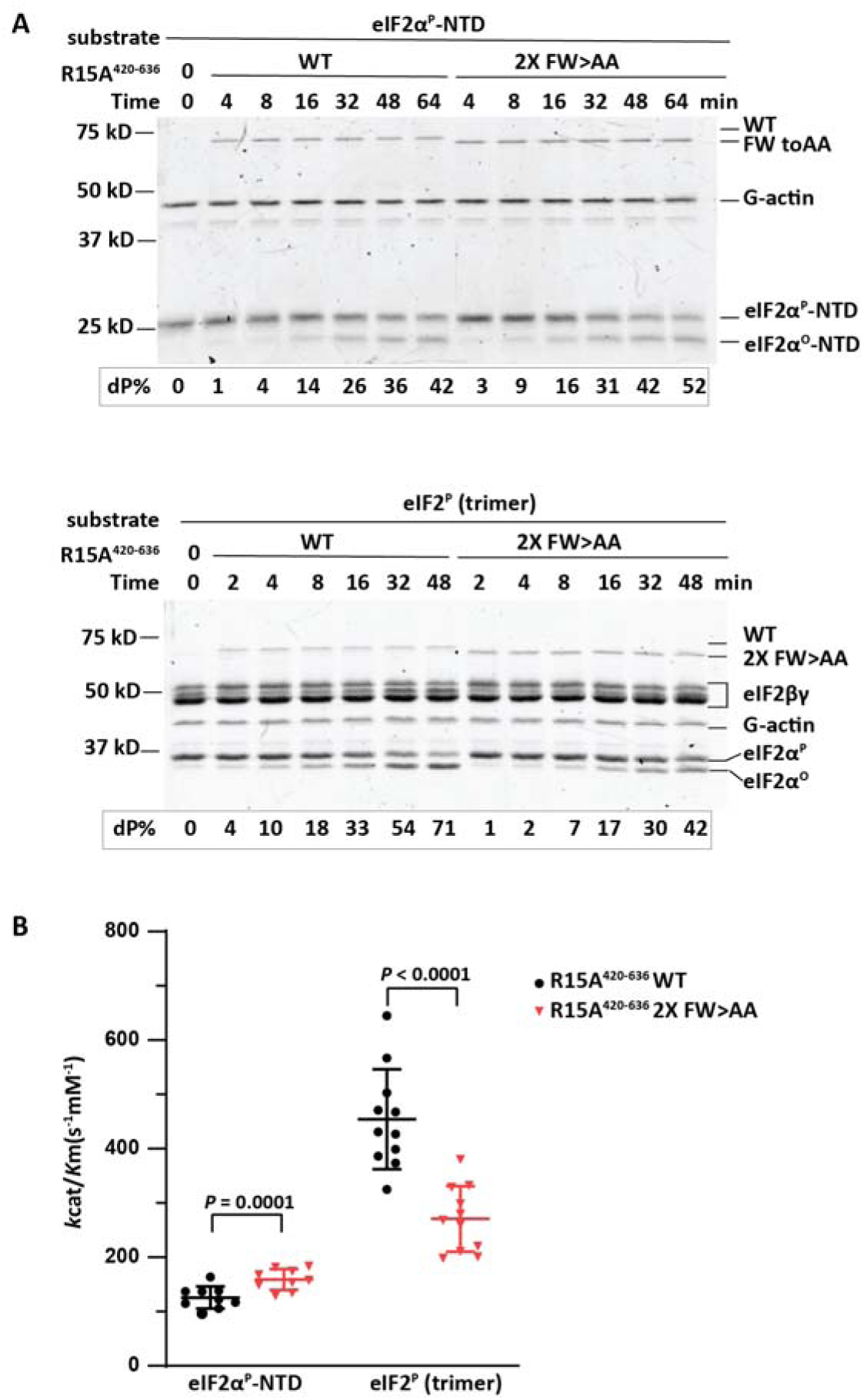
The eIF2γ-contacting Phe and Trp residues of PPP1R15A contribute to eIF2^P^ dephosphorylation in vitro, by reduced PPP1R15A^420-636^-containing holophosphatase confined to repeats 3 and 4. (A) Coomassie stained SDS-PAGE PhosTag gels of dephosphorylation reactions of the core substrate (eIF2α^P^_NTD, upper panel) or the physiological substrate (eIF2^P^ trimer, lower panel) by a holophosphatase comprised of the wildtype PPP1R15A^420-636^ (reduced to repeats 3 and 4) or its 2X FW>AA mutant counterpart (F428A, W432A, F479A, W482A), G-actin and PP1A. Shown is a representative of experiments reproduced three times. (B) Graphic display of the mean ± s.d. of the *k*_cat_/*K*_m_ extracted from all experimental points with the indicated substrates and holophosphatases. *P* values for two tailed parametric t-test are shown.

**Fig. S5.**
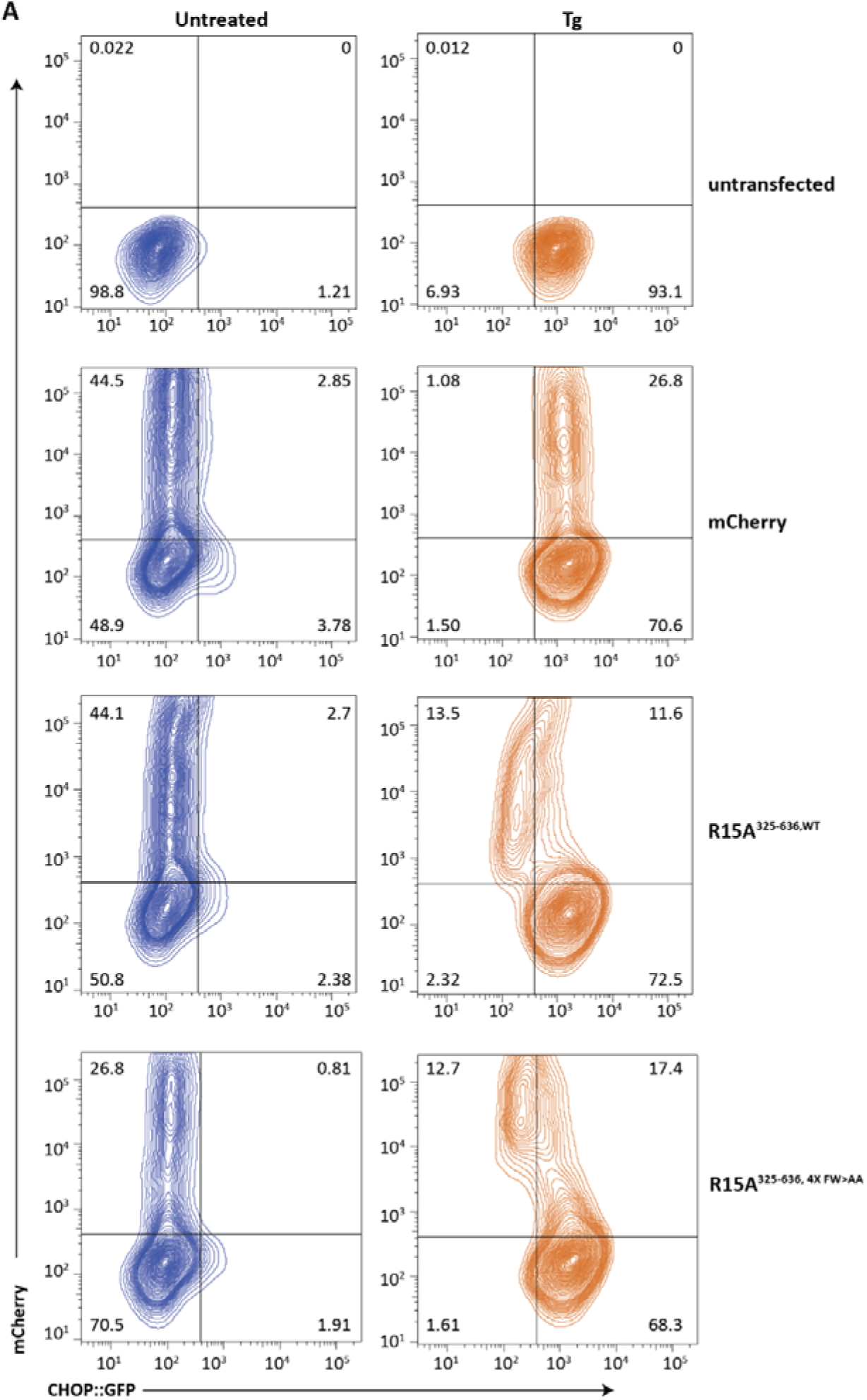
PPP1R15A contacts with eIF2γ contribute to ISR termination in cells transfected with mammalian expression plasmids encoding the counterpart to the PPP1R15A^325-636^ used in vitro. Dual channel flow cytometry plots of untreated and thapsigargin (Tg)-treated CHOP:GFP transgenic CHO-K1 cells transfected with mCherry alone or human PPP1R15A (wild type or mutants as indicated) with mCherry fused to their C-termini.

**Table S1:**
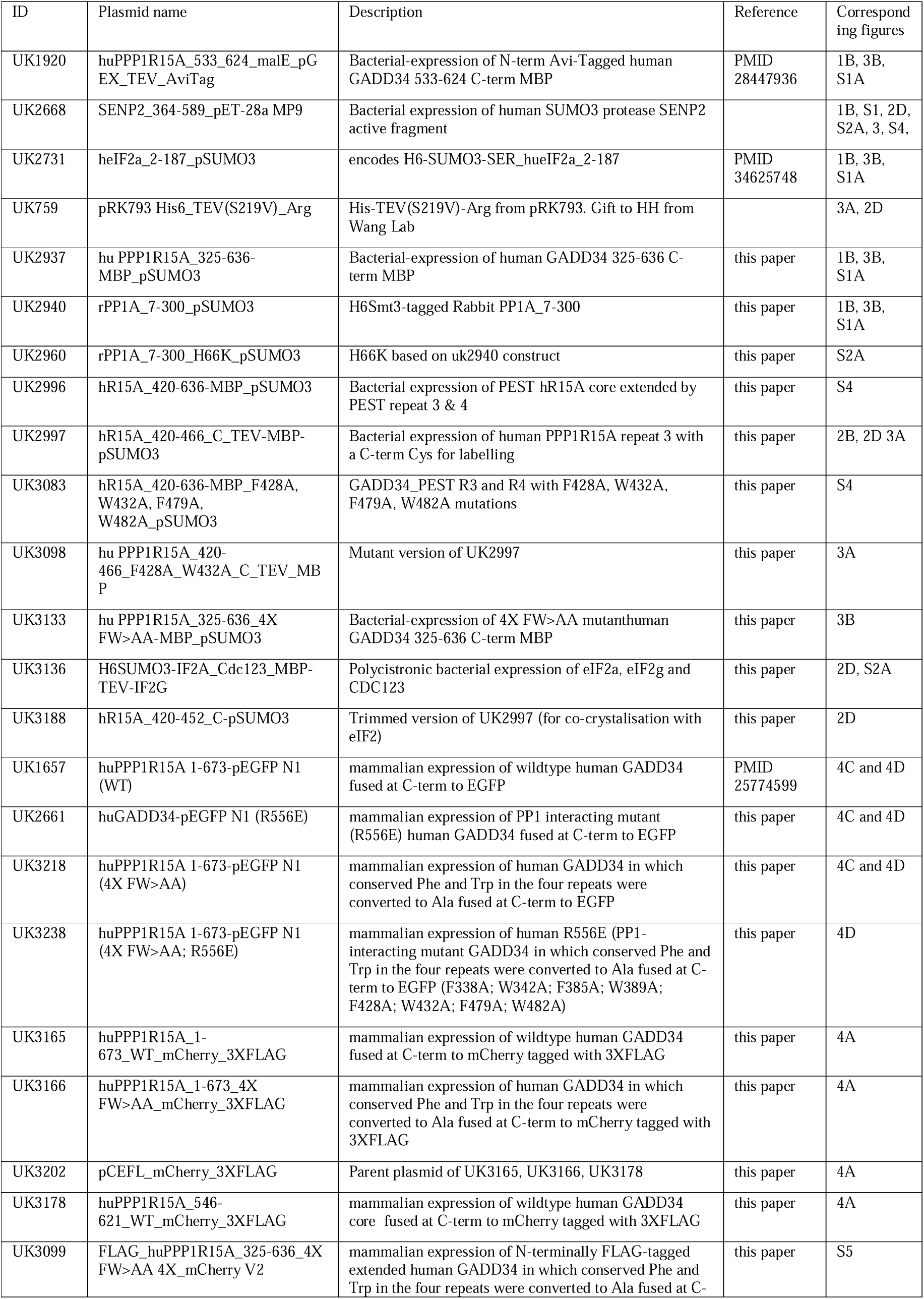
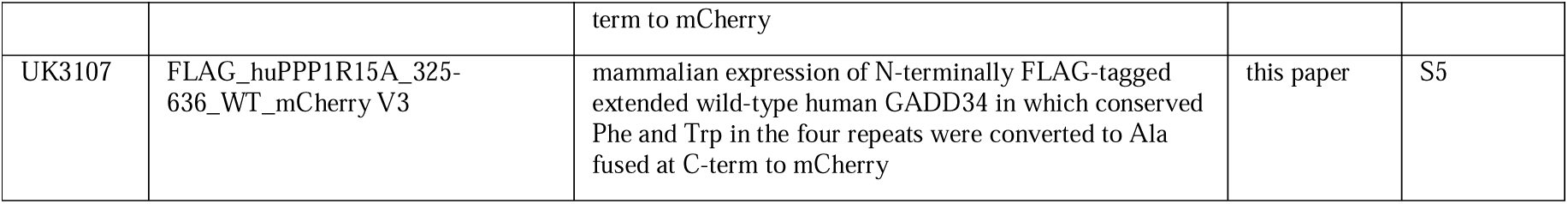
List and description of plasmids used in this study. ^a^ Values in parentheses are for the highest resolution shell.

**Table S2:**
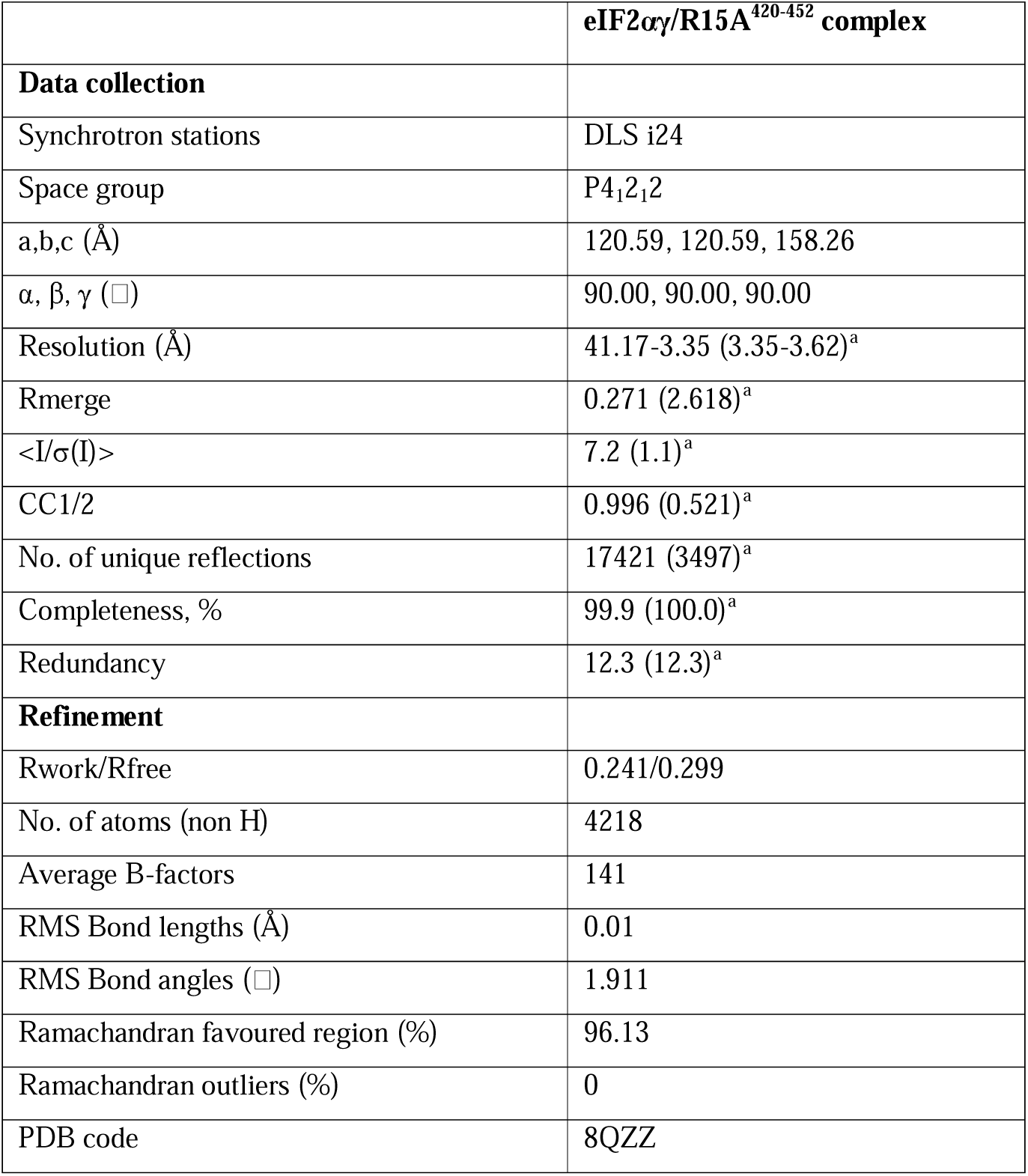
X-ray data collection and refinement statistics.

**Table S3.**
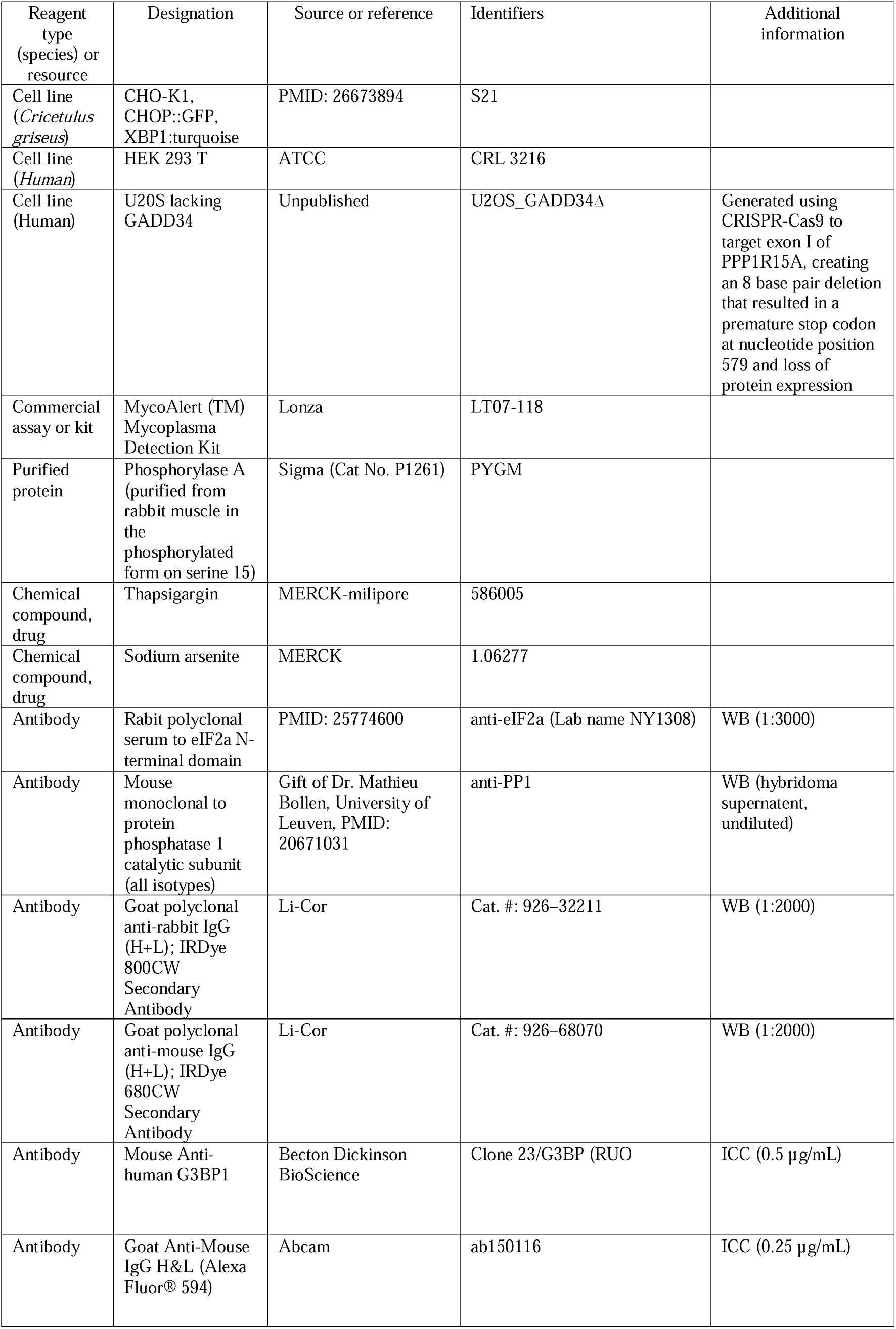
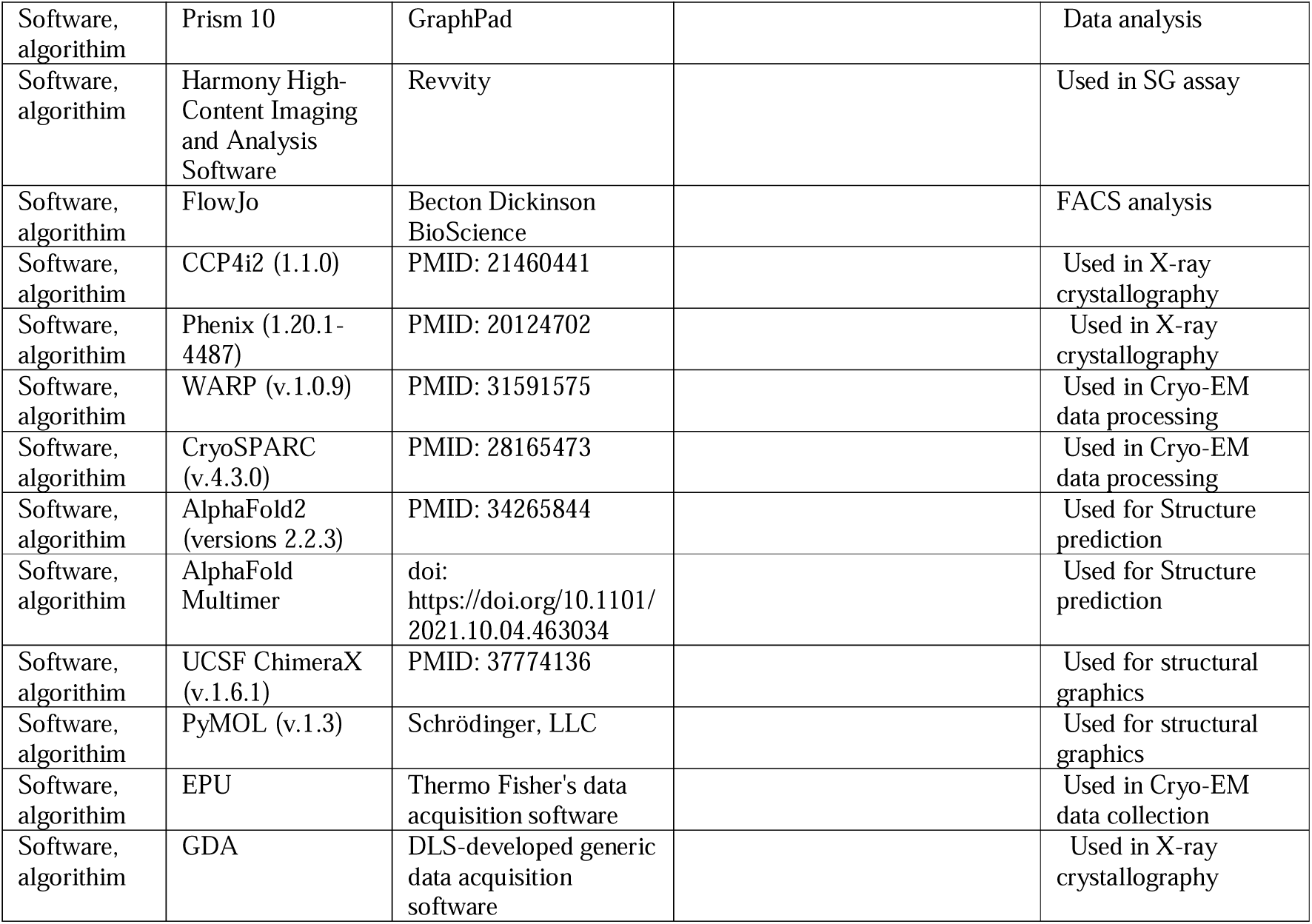
Key resource table.

## References

1. H. P. Harding et al., An integrated stress response regulates amino acid metabolism and resistance to oxidative stress. Mol Cell 11, 619–633 (2003).

2. A. G. Hinnebusch, I. P. Ivanov, N. Sonenberg, Translational control by 5’-untranslated regions of eukaryotic mRNAs. Science 352, 1413–1416 (2016).

3. M. Costa-Mattioli, P. Walter, The integrated stress response: From mechanism to disease. Science 368, eaat5314 (2020).

4. C. Jousse et al., Inhibition of a constitutive translation initiation factor 2alpha phosphatase, CReP, promotes survival of stressed cells. J Cell Biol 163, 767–775 (2003).

5. I. Novoa, H. Zeng, H. Harding, D. Ron, Feedback inhibition of the unfolded protein response by GADD34-mediated dephosphorylation of eIF2a. J Cell Biol 153, 1011–1022 (2001).

6. Y. Ma, L. M. Hendershot, Delineation of a Negative Feedback Regulatory Loop That Controls Protein Translation during Endoplasmic Reticulum Stress. J Biol Chem 278, 34864–34873 (2003).

7. I. Novoa et al., Stress-induced gene expression requires programmed recovery from translational repression. Embo J 22, 1180–1187 (2003).

8. E. Malzer et al., Coordinate regulation of eIF2alpha phosphorylation by PPP1R15 and GCN2 is required during Drosophila development. J Cell Sci 126, 1406–1415 (2013).

9. M. S. Choy et al., Structural and Functional Analysis of the GADD34:PP1 eIF2alpha Phosphatase. Cell reports 11, 1885–1891 (2015).

10. J. E. Chambers et al., Actin dynamics tune the integrated stress response by regulating eukaryotic initiation factor 2alpha dephosphorylation. eLife 4 (2015).

11. M. H. Brush, D. C. Weiser, S. Shenolikar, Growth arrest and DNA damage-inducible protein GADD34 targets protein phosphatase 1 alpha to the endoplasmic reticulum and promotes dephosphorylation of the alpha subunit of eukaryotic translation initiation factor 2. Mol Cell Biol 23, 1292–1303 (2003).

12. H. P. Harding et al., Ppp1r15 gene knockout reveals an essential role for translation initiation factor 2 alpha (eIF2alpha) dephosphorylation in mammalian development. Proc Natl Acad Sci U S A 106, 1832–1837 (2009).

13. R. Chen et al., G-actin provides substrate-specificity to eukaryotic initiation factor 2-alpha holophosphatases. eLife 4, e04871 (2015).

14. Y. Yan, H. P. Harding, D. Ron, Higher-order phosphatase-substrate contacts terminate the integrated stress response. Nat Struct Mol Biol 28, 835–846 (2021).

15. M. Rojas, G. Vasconcelos, T. E. Dever, An eIF2a-binding motif in protein phosphatase 1 subunit GADD34 and its viral orthologs is required to promote dephosphorylation of eIF2a. Proc Natl Acad Sci U S A 112, E3466–3475 (2015).

16. T. Ito, A. Marintchev, G. Wagner, Solution structure of human initiation factor eIF2alpha reveals homology to the elongation factor eEF1B. Structure 12, 1693–1704 (2004).

17. J. Brito Querido et al., Structure of a human 48S translational initiation complex. Science 369, 1220–1227 (2020).

18. M. Rojas, A. C. Gingras, T. E. Dever, Protein phosphatase PP1/GLC7 interaction domain in yeast eIF2gamma bypasses targeting subunit requirement for eIF2alpha dephosphorylation. Proc Natl Acad Sci U S A 111, E1344–1353 (2014).

19. N. L. Kedersha, M. Gupta, W. Li, I. Miller, P. Anderson, RNA-binding Proteins TIA-1 and TIAR Link the Phosphorylation of eIF-2alpha to the Assembly of Mammalian Stress Granules. J Cell Biol 147, 1431–1442 (1999).

20. C. Sidrauski, A. M. McGeachy, N. T. Ingolia, P. Walter, The small molecule ISRIB reverses the effects of eIF2alpha phosphorylation on translation and stress granule assembly. eLife 4 (2015).

21. A. Ruggieri et al., Dynamic oscillation of translation and stress granule formation mark the cellular response to virus infection. Cell Host Microbe 12, 71–85 (2012).

22. J. Chou, B. Roizman, The gamma 1(34.5) gene of herpes simplex virus 1 precludes neuroblastoma cells from triggering total shutoff of protein synthesis characteristic of programed cell death in neuronal cells. Proc Natl Acad Sci U S A 89, 3266–3270 (1992).

23. B. He, M. Gross, B. Roizman, The gamma(1) 34.5 protein of herpes simplex virus 1 has the structural and functional attributes of a protein phosphatase 1 regulatory subunit and is present in a high molecular weight complex with the enzyme in infected cells. J Biol Chem 273, 20737–20743 (1998).

24. W. Shi et al., GADD34-PP1c recruited by Smad7 dephosphorylates TGF{beta} type I receptor. J Cell Biol 164, 291–300 (2004).

25. S. J. Marciniak et al., CHOP induces death by promoting protein synthesis and oxidation in the stressed endoplasmic reticulum. Genes and Development 18, 3066–3077 (2004).

26. M. D’Antonio et al., Resetting translational homeostasis restores myelination in Charcot-Marie-Tooth disease type 1B mice. J Exp Med 210, 821–838 (2013).

27. I. Das et al., Preventing proteostasis diseases by selective inhibition of a phosphatase regulatory subunit. Science 348, 239–242 (2015).

28. M. Carrara, A. Sigurdardottir, A. Bertolotti, Decoding the selectivity of eIF2alpha holophosphatases and PPP1R15A inhibitors. Nat Struct Mol Biol 24, 708–716 (2017).

29. A. Crespillo-Casado, J. E. Chambers, P. M. Fischer, S. J. Marciniak, D. Ron, PPP1R15A-mediated dephosphorylation of eIF2alpha is unaffected by Sephin1 or Guanabenz. eLife 6, e26109 (2017).

30. A. Crespillo-Casado et al., A Sephin1-insensitive tripartite holophosphatase dephosphorylates translation initiation factor 2alpha. J Biol Chem 293, 7766–7776 (2018).

31. Y. L. Wong et al., The small molecule ISRIB rescues the stability and activity of Vanishing White Matter Disease eIF2B mutant complexes. eLife 7 (2018).

32. G. Winter et al., DIALS: implementation and evaluation of a new integration package. Acta Crystallogr D Struct Biol 74, 85–97 (2018).

33. P. R. Evans, G. N. Murshudov, How good are my data and what is the resolution? Acta Crystallogr D Biol Crystallogr 69, 1204–1214 (2013).

34. M. D. Winn et al., Overview of the CCP4 suite and current developments. Acta Crystallogr D Biol Crystallogr 67, 235–242 (2011).

35. A. J. McCoy et al., Phaser crystallographic software. J Appl Crystallogr 40, 658–674 (2007).

36. P. D. Adams et al., PHENIX: a comprehensive Python-based system for macromolecular structure solution. Acta Crystallogr D Biol Crystallogr 66, 213–221 (2010).

37. P. Emsley, B. Lohkamp, W. G. Scott, K. Cowtan, Features and development of Coot. Acta Crystallogr D Biol Crystallogr 66, 486–501 (2010).

38. G. N. Murshudov et al., REFMAC5 for the refinement of macromolecular crystal structures. Acta Crystallogr D Biol Crystallogr 67, 355–367 (2011).

39. P. V. Afonine et al., Real-space refinement in PHENIX for cryo-EM and crystallography. Acta Crystallogr D Struct Biol 74, 531–544 (2018).

40. V. B. Chen et al., MolProbity: all-atom structure validation for macromolecular crystallography. Acta Crystallogr D Biol Crystallogr 66, 12–21 (2010).

41. E. C. Meng et al., UCSF ChimeraX: Tools for Structure Building and Analysis. Protein Sci 10.1002/pro.4792, e4792 (2023).

42. D. Tegunov, P. Cramer, Real-time cryo-electron microscopy data preprocessing with Warp. Nat Methods 16, 1146–1152 (2019).

43. A. Punjani, J. L. Rubinstein, D. J. Fleet, M. A. Brubaker, cryoSPARC: algorithms for rapid unsupervised cryo-EM structure determination. Nat Methods 14, 290–296 (2017).

44. E. Richard et al., Protein complex prediction with AlphaFold-Multimer. bioRxiv 10.1101/2021.10.04.463034, 2021.2010.2004.463034 (2022).

45. J. Jumper et al., Highly accurate protein structure prediction with AlphaFold. Nature 596, 583–589 (2021).

46. M. J. Abraham et al., GROMACS: High performance molecular simulations through multi-level parallelism from laptops to supercomputers. SoftwareX 1-2, 19–25 (2015).

47. Y. Yu et al., Semi-automated Optimization of the CHARMM36 Lipid Force Field to Include Explicit Treatment of Long-Range Dispersion. J Chem Theory Comput 17, 1562–1580 (2021).

48. G. Bussi, D. Donadio, M. Parrinello, Canonical sampling through velocity rescaling. J Chem Phys 126, 014101 (2007).

49. M. Parrinello, A. Rahman, Polymorphic transitions in single crystals: A new molecular dynamics method. Journal of Applied Physics 52, 7182–7190 (1981).

50. U. Essmann et al., A smooth particle mesh Ewald method. The Journal of Chemical Physics 103, 8577–8593 (1995).

51. S. Miyamoto, P. A. Kollman, Settle: An analytical version of the SHAKE and RATTLE algorithm for rigid water models. Journal of Computational Chemistry 13, 952–962 (1992).

52. B. Hess, P-LINCS:L A Parallel Linear Constraint Solver for Molecular Simulation. Journal of Chemical Theory and Computation 4, 116–122 (2008).

53. W. Humphrey, A. Dalke, K. Schulten, VMD: visual molecular dynamics. J Mol Graph 14, 33-38, 27–38 (1996).

54. E. F. Pettersen et al., UCSF Chimera--a visualization system for exploratory research and analysis. J Comput Chem 25, 1605–1612 (2004).

55. M. Golicnik, Explicit analytic approximations for time-dependent solutions of the generalized integrated Michaelis-Menten equation. Anal Biochem 411, 303–305 (2011).

56. N. Van Dessel et al., The phosphatase interactor NIPP1 regulates the occupancy of the histone methyltransferase EZH2 at Polycomb targets. Nucleic Acids Res 38, 7500–7512 (2010).

